# B cells discriminate HIV-1 Envelope protein affinities by sensing antigen binding association rates

**DOI:** 10.1101/2022.01.14.476215

**Authors:** Md. Alamgir Hossain, Kara Anasti, Brian Watts, Kenneth Cronin, Advaiti Pai Kane, R.J. Edwards, David Easterhoff, Jinsong Zhang, Wes Rountree, Yaneth Ortiz, Laurent Verkoczy, Michael Reth, S. Munir Alam

## Abstract

HIV-1 Envelope (Env) proteins designed to induce neutralizing antibody responses allow study of the role of affinities (equilibrium dissociation constant, K_D_) and kinetic rates (association/dissociation rates) on B cell antigen recognition. It is unclear whether affinity discrimination during B cell activation is based solely on Env protein binding K_D_, and whether B cells discriminate between proteins of similar affinities but that bind with different kinetic rates. Here we used a panel of Env proteins and Ramos B cell lines expressing IgM BCRs with specificity for CD4 binding-site broadly neutralizing (bnAb) or a precursor antibody to study the role of antigen binding kinetic rates on both early (proximal/distal signaling) and late events (BCR/antigen internalization) in B cell activation. Our results support a kinetic model for B cell activation in which Env protein affinity discrimination is based not on overall K_D_, but on sensing of association rate and a threshold antigen-BCR half-life.

## INTRODUCTION

The binding of cognate antigen to the IgM-class B cell antigen receptor (IgM-BCR) initiates signaling and B cell activation (Packard and Cambier, 2013; Reth and Wienands, 1997). However, the mechanism by which B cells sense antigen and initiate signaling leading to full activation remains unclear. Recent super-resolution microscopy studies oppose the classical view in which BCRs are monomeric, freely diffusing receptors that only cluster upon antigen binding (Pierce and Liu, 2010; Schamel and Reth, 2000; Tolar et al., 2009). Instead, on the surface of resting B cells the IgM-BCR forms closed, autoinhibited oligomers that reside in particular membrane compartments, and B cell activation is dependent on the ability of an antigen to dissociate the closed and presumably laterally shielded IgM-BCR so that it can gain access to coreceptors such as CD19 for signaling (Maity et al., 2015; Schamel and Reth, 2000; Yang and Reth, 2010a, b). Thus, with these revisions to the classical models, there is a need to revisit the role of antigen affinity and valency in activating B cells that leads to development of protective antibody responses against viral pathogens, like HIV-1.

For classical haptens and proteins, B cells are activated by a wide range of affinities (μM to low nM, 10^-6^ – 10^-10^ M), corresponding to the theoretical limits of diffusion and physiologically consequential antigen-BCR complex dwell time (Foote and Eisen, 1995, 2000). However, while antigens with affinities as weak as 100μM can activate naïve B-cells, this requirement is influenced by clonal competition, with higher affinity naïve B cells outcompeting lower affinity counterparts (Dal Porto et al., 2002; Shih et al., 2002). In addition, HIV-1 Envelope (Env) proteins (with equilibrium dissociation constant, K_D_ <1μM) designed to target bnAb precursor B cells that are either infrequent (Abbott et al., 2018) and/or which have lower BCR densities as a result of tolerance-mediated functional silencing (anergy) (Chen et al., 2013; McGuire et al., 2016; Saunders et al., 2019; Verkoczy et al., 2010; Williams et al., 2017), are more effective immunogens when in multimeric form (Dennison et al., 2011; Verkoczy et al., 2011; Zhang et al., 2016). The role for avidity is consistent with the reported requirement of multivalent interactions for triggering a “quantized number of BCRs (n= 10 -20)” to induce B cell activation (Dintzis et al., 1982; Sulzer and Perelson, 1997; Vogelstein et al., 1982), although when the antigen is appropriately spaced, the minimal valency needed to induce signaling can be much lower (Veneziano et al., 2020).

From the above studies, it would thus appear that Env proteins with lower K_D_ values (higher affinities) would be superior HIV vaccine immunogens. In reality however, the predictive value of proteins having lower K_D_ for selecting immunogen candidates is unclear, since it has been experimentally demonstrated that antigens with excessively high affinities can prematurely prime naïve B cells to terminally differentiate (Chan and Brink, 2012; Paus et al., 2006; Phan et al., 2006). Furthermore, it has also been elegantly demonstrated that germinal center (GC) responses to complex antigens comprise a range of affinities in both early and late GC responses (Kuraoka et al., 2016), and GC+ B cells can be driven by considerably weaker affinity (K_D_ =40μM) protein antigens (Dosenovic et al., 2018) that impact decisions on B cell affinity maturation and fate (to either memory B-cells or terminally-differentiated plasmacytes) (Paus et al., 2006; Viant et al., 2020; Viant et al., 2021). The above studies also imply that avidity could be a major discriminator of Env immunogenicity. However, it is unlikely that B cells rely solely on avidity for affinity discrimination, since B cells with weaker affinity BCRs would benefit far more than those with higher-affinity, and a graded response to affinity would not occur.

One key caveat in the majority of the above studies is that the affinity read-outs were based on equilibrium dissociation constant (K_D_) and therefore, either attention to association (k_a_) or dissociation (k_d_) rate constant was not made or the immunogen design process resulted in the selection of protein antigens that bind with similarly fast k_a_ values. The role of dissociation kinetics in antigen presentation in a study only compared antigens with similar k_a_ values approaching the diffusion limit (Batista and Neuberger, 1998); thus, the role of association rate was not studied. In anti-HIV-1 responses, affinity maturation of bnAbs involves sequential improvement in both k_a_ and k_d_, with improvement in k_a_ often preceding that of k_d_ (Bonsignori et al., 2012; Henderson et al., 2019). The observed k_a_ improvement early in bnAb development suggests that an optimal association interaction likely affords a selection advantage (Henderson et al., 2019). It is unlikely that antigen binding to the BCRs on a cognate B cell reaches an equilibrium state *in-vivo* and therefore, it raises the question whether design and down-selection of immunogens should be strictly based on K_D_ values. Given the different temporal windows during which B cell signaling (1-2 min) and BCR/antigen internalization (∼30 min) occurs, we hypothesize that B cell activation is dependent on the kinetic rates and requires an affinity with optimal k_a_/k_d_ rates.

We recently designed HIV Env proteins that have different affinities to well-defined affinity-matured and inferred germline antibodies (Saunders et al., 2017; Saunders et al., 2019), and which therefore present a unique test set for elucidating how HIV-1 Env proteins are sensed by naïve B cells to initiate early signaling events that lead to BCR/antigen internalization. Because these early events are key steps to peptide presentation on MHC class II and subsequent recruitment of T cell help, understanding these initial steps using well-defined in vitro, ex vivo, and in vivo B-cell systems is crucial. Here, we developed Ramos B cell lines that expressed IgM BCRs with specificities of HIV-1 CD4 binding-site bnAbs (CH31 or VRC01) or the VRC01 unmutated common ancestor (UCA) antibodies (Bonsignori et al., 2012; Bonsignori et al., 2018). Using a panel of Env proteins that show binding to the CD4 binding-site antibodies with varying affinities and kinetic rates, we studied B cell proximal and distal signaling as well as BCR/antigen internalization to determine how antigen binding rates influence B cell activation and the BCR endocytic function in antigen internalization. We report that the strength of B cell activation is not dependent on the dissociation equilibrium constant (K_D_) but on the association rate that impact phospho-signaling kinetics leading to calcium mobilization. In addition, our studies show that BCR-antigen internalization has a threshold and a ceiling that are dependent on both the association rate and the half-life of the antigen-bound BCR complex, indicating a required minimum threshold for antigen binding dwell time for BCR-antigen internalization. Our studies provide an explanation of how B cells sense affinities to complex protein antigens and the importance of designing HIV-1 immunogens with the optimal Env-antibody binding association rate and an above threshold BCR-antigen dissociation half-life.

## RESULTS

### HIV-1 Env proteins with varying affinities and kinetic rates

For this study, we selected the CH31 bnAb (Bonsignori et al., 2012), a member of a clone of five V_H_1-2-utilizing, CD4bs-directed VRC01-class bnAbs (CH30-34), which underwent somatic hypermutation (SHM) resulting in 24% V_H_DJ_H_ nucleotide changes, and a net 9 aa insertion (6-nt deletion and a 33-nt tandem duplication insertion) event in the HCDR1 (Kepler et al., 2014). Previously, we had reported that the inclusion of the indel duplicon in CH31 UCA mutant antibody resulted in an 8-fold increase in the k_a_, and conversely, CH31 intermediates that lacked this indel bound with a k_a_ slower by one order of magnitude, indicating the importance of association kinetic rate improvement during the early stage of bnAb affinity maturation (Henderson et al., 2019; Kepler et al., 2014). To select antigens with varying affinities and kinetics rates, we used a panel of HIV-1 Envelope (Env) proteins, each in either monomeric or trimeric form, and measured the affinities and kinetics rates of binding (association rate, k_a_ and dissociation rate, k_d_) to CH31 IgG. The Env proteins included germline-targeting (GT) monomeric forms of outer domain of gp120 (Jardine et al., 2013), 426c core proteins derived from the clade C 426c Env (Bonsignori et al., 2017; McGuire et al., 2013), monomeric gp120 proteins, and trimeric gp140 proteins (Liao et al., 2013; Saunders et al., 2017; Saunders et al., 2019) (**Fig. 1**). The GT monomers, GT8 and GT6, designed to bind to the CD4bs bnAb VRC01 precursor antibodies with high affinities (Jardine et al., 2013), demonstrated fast association (k_a_ >1x10^4^ M^-1^s^-1^) and dissociation rates (k_d_ >1x10^-2^s^-1^) to CH31 (**Fig. 1C**). GT8, with both relatively faster k_a_ and slower k_d_, demonstrated a higher overall affinity (K_D_ = 307.8nM) when compared to GT6 (K_D_ = 5.2μM) (**Fig. 1C**). 426c (K_D_ = 108.8 nM) had a much higher affinity to CH31 when compared to the deglycosylated form, 426c degly3 (K_D_ = 26.4μM), predominately due to its much slower dissociation rate (k_d_ = 3.0x10^-4^ s^-1^ versus 7.4x10^-1^ s^-1^) (**Fig. 1C**), indicating that glycan site modifications that gave VRC01 germline reverted antibody binding (McGuire et al., 2013) impact binding of CH31 bnAb. Among the monomeric gp120 proteins, CH505TF gp120 bound with faster kinetic rates to CH31 than A244 gp120 (**Fig. 1C**). As a result of its much slower dissociation rate (k_d_<1x10^-4^ s^-1^), the overall affinity of A244 gp120 to CH31 was approximately 10-fold higher than CH505TF gp120 (**Fig. 1C**). While the gp140 trimers bound to CH31 with moderate to high affinities (K_D_ = 56 – 2nM), binding of the trimers displayed kinetic rates that were orders of magnitude different from each other (**Fig. 1B-C**). Thus, the selected panel of monomeric and trimeric Env proteins provided a wide range of affinities (10^-5^ to 10^-9^ M) and kinetic rates allowing us to determine whether or not B cells sense affinity solely based on the K_D_ values (**Fig. 1**).

**Figure 1.**
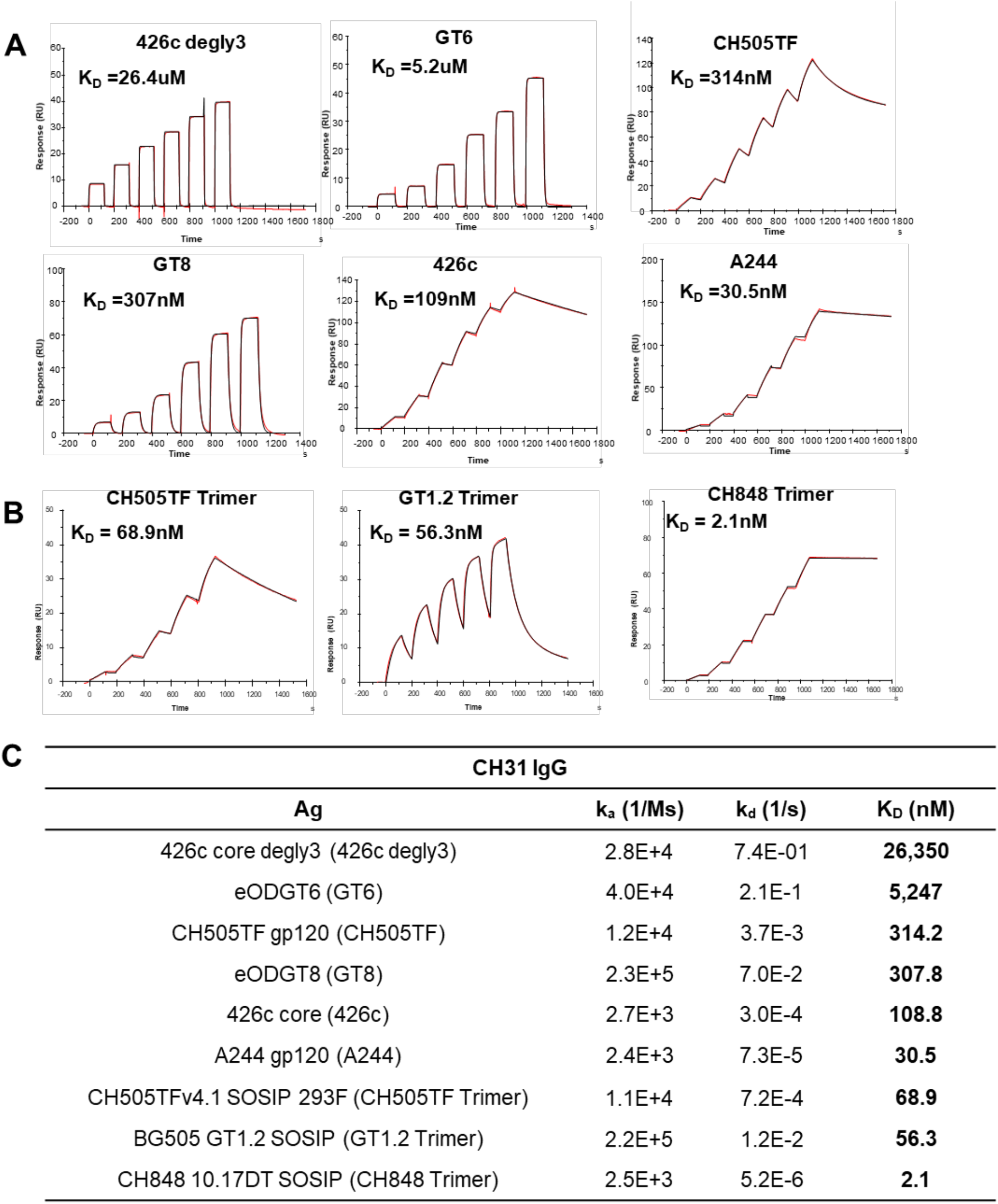
Affinities of Env proteins to CH31. (**A**) Surface Plasmon Resonance (SPR) single-cycle kinetic binding profiles and affinities of Env proteins (core and gp120) to the CD4 binding site bnAb CH31 IgG. Six sequential injections of each antigen at concentrations ranging from 25nM to 5000nM were flowed over CH31 IgG mAb. Curve fitting analyses were performed using either a 1:1 Langmuir model or the heterogeneous ligand model. (**B**) Single-cycle kinetic binding profiles and affinities of CH31 IgG Fab (30-1500nM) against biotinylated GT1.2, CH505TF, or CH848 trimers immobilized to streptavidin coated sensor chip. The heterogeneous ligand model was used for fitting CH31 Fab against GT1.2 Trimer and the 1:1 Langmuir model was used for fitting against both the CH505TF and CH848 Trimers. (**C**) Binding affinities (K_D_) and kinetic rate constants (association and dissociation rates) of each protein to either CH31 IgG mAb or Fab. Binding curves and values for the kinetic rates and affinities are representative of at least two independent measurements.

### B cell signaling is dependent on Env protein-binding association rate

To measure the strength of B cell activation by antigens of differing affinities, a Ramos cell line was developed expressing the CH31 derived IgM^+^ BCRs (CH31 IgM cells). Although affinity and kinetic rates measurements were done using IgG-class antibodies (**Fig. 1**), a comparison of CH31 IgG and CH31 IgM antibodies showed that the two isoforms maintain not only the same specificity but also bind with similar affinities to the tested proteins (**Fig. S1**). Phenotypic analysis of cell surface markers on CH31 IgM cells showed expression of IgM and the relevant κ-light chain, as well as key coreceptor and accessory molecules (**Fig. S2A**). Exposure of the CH31 IgM cells to anti-IgM Fab_2_ resulted in a strong calcium flux (Ca-flux) response indicating that the CH31 IgM-BCR on Ramos cells is functional (**Fig. S2B**).

We first measured Ca-flux responses, an early indicator of B cell activation, to three trimeric gp140 proteins (**Fig. 2A**). Among these proteins, GT1.2 is a next generation BG505 SOSIP trimer designed to target the VRC01-like bnAb lineages (Sanders et al., 2015), including CH31, while CH505TF and CH848 trimers were developed to target the CD4bs (‘HCDR3 binder’-specific) subclass CH103 bnAb lineage (Liao et al., 2013) and the V3-glycan-directed DH270 bnAb lineage (Saunders et al., 2019) respectively. Negative-stain EM (NS-EM) analysis showed that the apparent CD4bs epitope, stoichiometry (n=3) and the angle of approach for the CH31 Fab were all similar for each of the three trimers (**Fig. S3**). However, examination of CH31 binding rates to the three gp140 trimers revealed varying affinities and kinetic rates. Specifically, two trimers, CH505TF and GT1.2, bound with similar affinities (68.9 and 56.3 nM), while CH848 trimer bound with an order of magnitude higher affinity (2.1nM) (**Fig. 1C**). The relatively weaker affinity GT1.2 trimer bound CH31 with the fastest association-rate (k_a_ = 2.2x10^5^ M^-1^s^-1^), almost two orders of magnitude faster than the high-affinity CH848 trimer that bound with the slowest k_a_ (k_a_ = 2.5x10^3^ M^-1^s^-1^) (**Fig. 1C**). In contrast, k_d_ decreased by orders of magnitude with increasing trimer affinities (**Fig. 1C**). Thermodynamics analysis showed entropy-enthalpy compensation during CH31 binding to the two trimers (GT1.2, CH505TF) (**Fig. S4, S5).** When compared to CH505TF, GT1.2 bound with a relatively lower entropic hurdle (-TΔS) and a negligible ΔCp value, a measure of conformational change during binding. In contrast, the binding of CH31 to the high-affinity CH848 trimer was associated with a much larger ΔCp value and a thermodynamically favorable -TΔS value, indicating distinct dynamics and an entropically-driven formation of the bound complex (**Fig. S5**). Overall, the above results showed that while CH31 engaged the same CD4bs epitope on the three trimers, the kinetic rates and the thermodynamic mechanisms of the interactions were markedly different.

**Figure 2.**
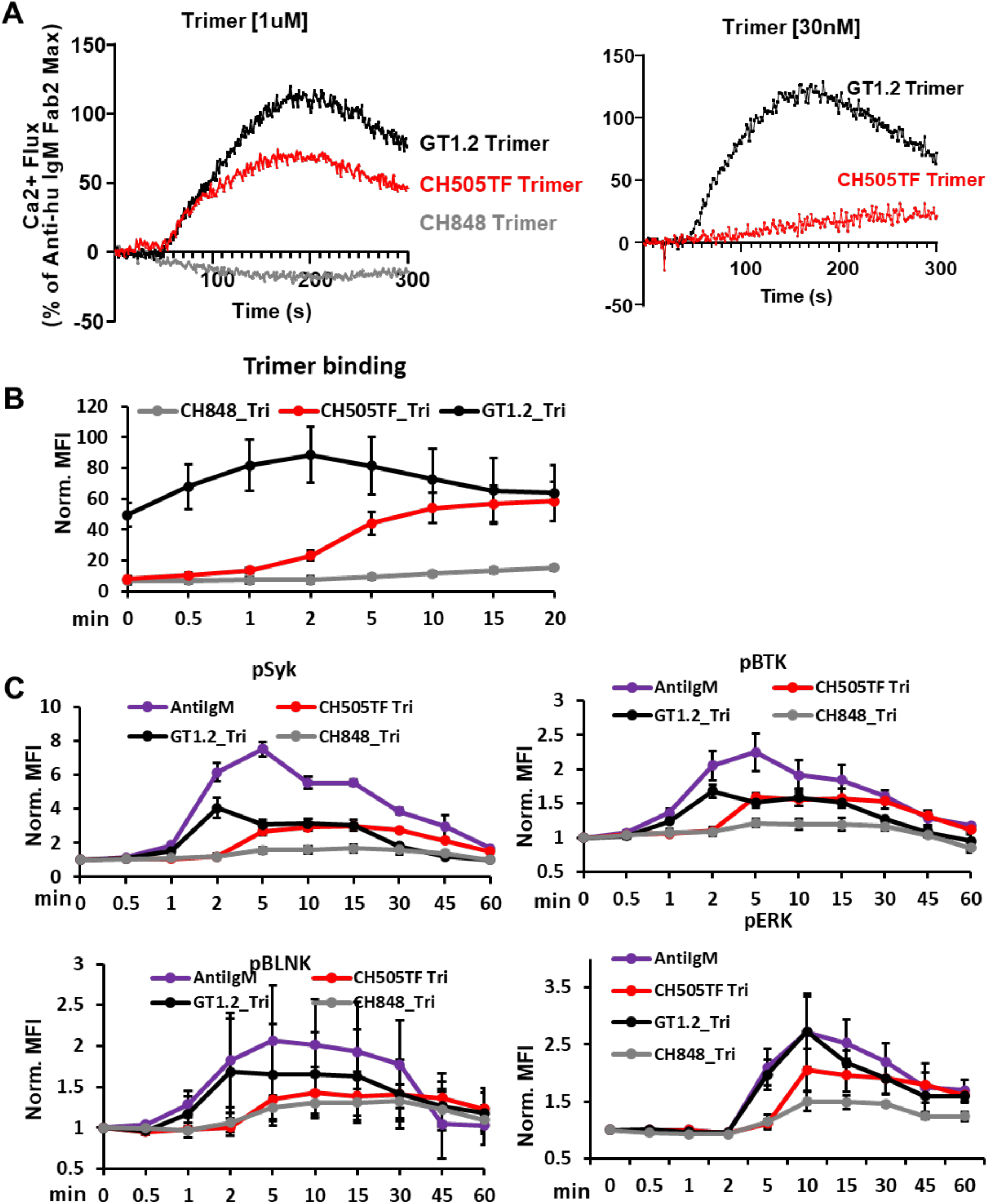
Signaling induced in Ramos cells expressing CH31 IgM by Env trimers. (**A**) Calcium flux responses induced in CH31 IgM Ramos cells following exposure to Env trimer proteins. Overlaid responses shown are to GT1.2 (black), CH505TF (red), or CH848 (gray) trimers at two different concentrations, 1μM (left) and 30nM (right). Calcium flux results are presented as a % of the maximum anti-human IgM F(ab)2 response and are representative of at least two measurements. (**B**) Time-course of binding of trimeric Env proteins to CH31 IgM cells. Ramos cells expressing CH31 IgM BCRs were incubated with biotinylated GT1.2 (black), CH505TF (red), or CH848 (gray) trimer along with anti-human IgM (purple), each at a concentration of 30nM. Ramos cells were collected at each time point, fixed and stained with anti-biotin PE antibody to detect binding of trimeric proteins over time. Median fluorescent intensity (MFI) value obtained from flow data analysis of treated samples were normalized to MFI value of untreated samples. Data plotted show mean values of the normalized MFI (Norm. MFI) measured from three independent experiments, and error bars represent the standard deviation. **(C)** Cell signaling kinetics following stimulation by trimeric Env proteins. The kinetics of Syk, BLNK, Btk or ERK1/2 phosphorylation over 1 h were measured in CH31 IgM Ramos cells exposed to 30 nM of either GT1.2 (black), CH505TF (red), or CH848 (gray) trimers along with anti-human IgM antibody (purple). Cells were pooled at each indicated time, fixed, permeabilized and subsequently stained to detect intracellular level of phosphorylated Syk, BLNK, Btk and ERK1/2. MFI values obtained for each treatment point were normalized to MFI values of unstimulated sample. Data plotted in the graphs are mean values of the normalized MFI with standard deviation calculated from three independent experiments.

Surprisingly, the high-affinity CH848 trimer failed to induce Ca-flux, while the relatively weaker-affinity trimers GT1.2 and CH505TF trimers triggered strong responses (**Fig. 2A**). When GT1.2 and CH505TF were further tested at a lower concentration that was closer to the binding K_D_ values (30nM), GT1.2 (with the fastest k_a_ = 2.2x10^5^ M^-1^s^-^1) induced a strong Ca-flux response, while CH505TF (with a relatively slower k_a_ = 1.1x10^4^ M^-1^s^-^1) induced a drastically reduced response (**Fig. 2A**). Thus, the ability to trigger Ca-flux and the magnitude of the response was determined by the trimer’s binding association rate and not its affinity (K_D_). The lack of induction of Ca-flux by the high-affinity trimer was due to its very weak binding to BCRs on CH31-IgM cells (**Fig. 2B**). Furthermore, the kinetics of each of the trimer surface interactions with CH31 IgM cells mirrored the differences in their SPR-determined k_a_ values (**Fig. 2B, S6A**). Specifically, those trimers with relatively slower k_a_ (CH505TF, CH848), exhibited very weak surface binding after 2 min of antigen exposure (when peak Ca-flux was detected with GT1.2) (**Fig. 2B**). Notably, CH848 (the trimer with the slowest k_a_), maintained low surface binding, even after 20 min of antigen exposure. Thus, these trimer-BCR surface binding profiles demonstrate that Ca-mobilization is strongly dependent on antigen association kinetics and indicate a requirement of a BCR occupancy threshold, during the short (1-2 min) temporal window following antigen exposure.

To further evaluate the impact of differences in antigen binding kinetic rates on B cell signaling, we measured antigen-binding induced phosphorylation of several key signaling components. We monitored the kinetics and magnitude of phosphorylation of three proximal signaling molecules, namely the protein tyrosine kinases Syk and Btk as well as the adaptor protein (BLNK/SLP-65), that comprise components of the upstream signal transducing complex and mediate calcium mobilization via activation of phospholipase Cψ2 (PLCψ2) (Jumaa et al., 2005; Kurosaki, 2002; Lane et al., 1991; Takata et al., 1994). In addition, we measured phosphorylation of the extracellular signal-regulated kinase 1/2 (ERK1/2), a downstream signaling effector in the mitogen-activated protein kinases (MAPK) pathway that is associated with cell survival and proliferation (Jiang et al., 1998; Mizuno and Rothstein, 2005). Phosphorylation of each of the three proximal signaling molecules in response to the GT1.2 trimer was detected early (∼1min) and reached peak responses within 2 mins (**Fig. 2C**). In contrast, CH505TF trimer-induced phosphorylation of each of the signaling molecules exhibited slower kinetics (with responses peaking between 5-10 min) and was weaker in magnitude, particularly for BLNK and ERK (**Fig. 2C**). Despite its higher affinity, the markedly slower binding rate of the CH848 trimer (**Fig. 2B**) resulted in the weakest phosphorylation of both the proximal and distal signaling molecules (**Fig. 2C**), explaining its inability to induce Ca-mobilization (**Fig. 2A**).

To assess whether Ca-flux would be enhanced by multimerization of the three SOSIP trimers on nanoparticles (NP), we used GT1.2 SOSIP NP with 20 trimeric units (20-mer) as well as CH505TF and CH848 SOSIP NPs, each with 8 trimeric units (8-mers). The 8-mer of the high-affinity, slow association-rate binding trimer CH848 did not induce any Ca-flux (**Fig. S6B**). However, multimers of the SOSIP trimers, each binding with faster association rates (10^4^ - 10^5^ M^-1^s^-1^), did induce flux, but without any further enhancement in the Ca-mobilization magnitude relative to their standard trimeric forms (**Fig. S6B**). When compared to GT1.2 trimers, peak phosphorylation of the proximal signaling molecules, Syk and BLNK, in response to GT1.2 multimers (either the 20-mer NP described above, or additionally, made as a tetrameric NP) was slightly higher (<2-fold) (**Fig. S6C**), yet didn’t result in increased Ca-mobilization (**Fig. S6B**). Phosphorylation of either BTK or the distal Erk1/2 was also not enhanced by either of the GT1.2 multimers (**Fig. S6C**). Thus, the above results show that low valency trimeric interactions (binding stoichiometry n=3, **Fig. S3**) with faster association rates (k_a_ >10^4^ M^-1^s^-1^) are sufficient to trigger physiologically relevant phospho-signaling i.e. that which mediates strong Ca-mobilization. In contrast, the high affinity trimer (K_D_ = 2nM) with an order of magnitude slower association rate of binding (k_a_ ∼10^3^ M^-1^s^-1^) failed to induce Ca-flux, with or without multimerization. Hence, during activation of B cells expressing CD4bs bnAb IgM BCRs, avidity due to multivalent interactions cannot compensate for slower binding association rate, regardless of overall affinity (K_D_).

Since the above results indicate that the strength of B cell signaling is determined by Env trimer binding association-rate and not affinity (K_D_), this key assertion was further examined using an expanded panel of antigens that included several monomeric forms of the Env proteins (**Fig. 1**). In particular, tetrameric forms of the monomeric proteins were included and as such, their binding stoichiometry could be closely matched to that of the trimers (n=3, **Fig. S3**). SPR analysis of the affinities and kinetic rates for this panel of CH31-binding Env proteins showed that there was no association between their affinities (K_D_) or dissociation rates (k_d_, s^-1^) to Ca-flux (**Fig. 3A, 3B**). However, a positive and significant correlation was observed between the Env protein association rate (k_a_, M^-1^s^-1^) and Ca-flux (**Fig. 3C**, Kendall’s Tau =0.6000, p =0.0157). The proteins with the slowest association rates (k_a_< 3x10^3^ M^-1^s^-1^) triggered Ca-flux either weakly or not at all, whereas antigens with the fastest k_a_ (>10^4^ M^-1^s^-1^) mediated Ca-flux most potently, regardless of K_D_ value (**Fig. 3C**). Thus, in this expanded panel of CH31-binding Env immunogens, the strength of Ca-flux was associated with antigen binding association rate and not with antigen dissociation-rate or affinity, regardless of the antigen form (monomeric or trimeric).

**Figure 3.**
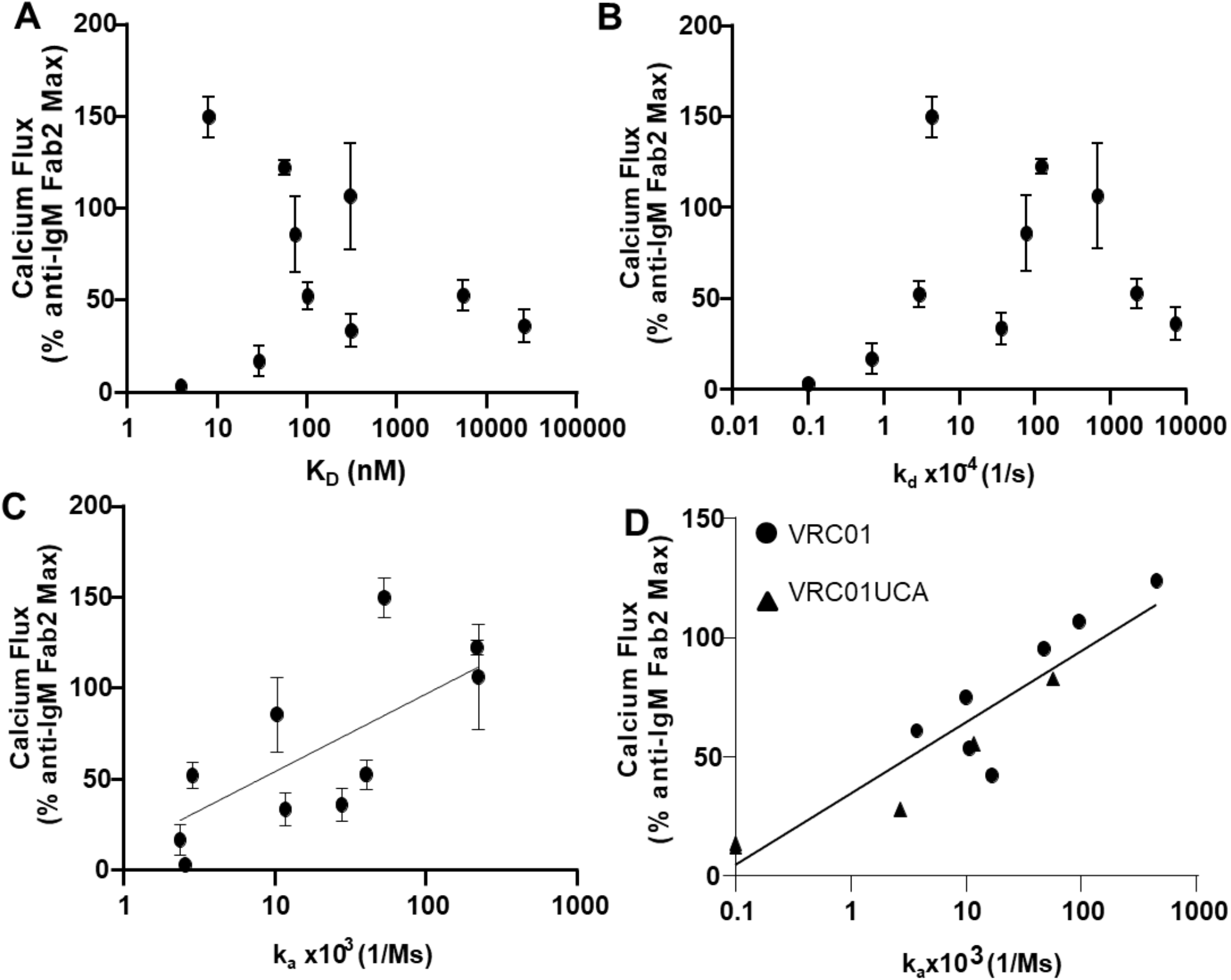
Relationship between calcium flux response and trimer binding kinetic rates. (**A**) CH31 IgM Ramos cell calcium flux induced by multimerized proteins (y-axis) versus SPR measured monomeric (or trimeric) protein affinity (K_D_, nM) to CH31 IgG (x-axis). (**B**) CH31 IgM Ramos cell calcium flux (y-axis) plotted versus monomeric antigen dissociation rates (k_d_, 1/s) with CH31 IgG bnAb (x-axis). (**C**) Calcium flux response (y-axis) versus protein association rates (k_a_, 1/Ms) to CH31 IgG (x-axis) with a semilog linear regression fit where x values are plotted on a logarithmic axis and y values are not (y=y-intercept+slope*Log(x)). (**D**) Calcium flux (y-axis) in VRC01 or VRC01UCA IgM Ramos cells plotted versus Env protein binding association rates (k_a_, 1/Ms) (x-axis) to CH31. VRC01 results are displayed as filled circles and VRC01UCA results are filled triangles. A semilog linear fit was applied to the VRC01/VRC01UCA data plotted in D. Calcium flux responses are plotted as a percentage of the maximum anti-human IgM F(ab)2 mediated responses and are the mean of at least two measurements with standard deviation error bars for CH31 and one measurement for VRC01 or VRC01UCA. Kinetic rate parameters and affinities are representative of at least two measurements for all IgGs. Correlation evaluation between k_a_ and Ca-flux was assessed by Kendall’s Tau analysis (Fig. C: Kendall’s Tau 0.6000; p-value=0.0157; Fig. D -VRC01 UCA: Kendall’s Tau 0.9487; p-value = 0.0230).

Finally, to examine if these results applied to other CD4bs-directed bnAbs, we tested Ramos cells expressing VRC01, as well as cells expressing its UCA Ab (Bonsignori et al., 2017) as IgM BCRs (herein referred to as VRC01 IgM and VRC01 UCA IgM cells, respectively; **Fig. S2**). Consistent with our above results using CH31 IgM cells, a positive relationship between the antigen association rates and Ca-flux was observed using either VRC01 or VRCO1 UCA IgM cells (**Fig. 3D, Fig. S7;** significant correlation for VRC01 UCA-IgM, Kendall’s Tau =0.9487, p =0.0230; not significant for VRC01-IgM). Specifically, only proteins that bound with k_a_ >10^3^ M^-1^s^-1^ induced detectable Ca-flux (>20% of reference Ab) (**Figure 3D**) and the magnitude of the response also increased with higher association rates, indicating that k_a_ above a threshold is required to initiate B cell activation. Together, these results show that the magnitude of the Ca-flux response is dependent on antigen binding association rate and not the dissociation rate.

### Role of antigen valency in B cell activation

To investigate the minimal antigen valency required for B cell activation, we compared the ability of monomeric versions of each of the Env proteins to induce Ca-flux in CH31 IgM cells, relative to their corresponding lower order (4-mer) multimers (formed following binding to the streptavidin molecule). None of the monomers tested mediated Ca-flux, but their tetrameric forms did, indicating that monomeric antigens require multimerization to activate CH31 IgM cells (**Fig. 4A**). Strikingly however, the magnitude of Ca-flux induced by these tetrameric proteins was not equal. In particular, tetrameric GT8 (with the fastest association rate, at k_a_>1x10^5^ M^-1^s^-1^) induced the strongest Ca-flux (**Fig. 4A**). On the other hand, tetrameric A244 gp120 (with the slowest binding k_a_=2.4x10^3^ M^-1^s^-1^ and strongest affinity), gave negligible Ca-flux responses. Furthermore, tetrameric forms of monomers with intermediate, relatively increasing binding association rates i.e., CH505TF gp120 (k_a_=1.2x10^4^ M^-1^s^-1^), 426c degly3 (k_a_=2.8x10^4^ M^-1^s^-1^) and GT6 (k_a_=4.0x10^4^ M^-1^s^-1^), mediated Ca-flux with increasing efficiency (**Fig. 4A**). Thus, while tetramerization of monomeric proteins was required to induce flux response, the magnitude of Ca-flux and phospho-signaling was proportional to their binding association rates (**Fig. 4B, S8**). Importantly, higher-order multimerization (exemplified by GT8 made as a 60-mer NP), did not show any enhancement of Ca-mobilization over that observed with the GT8-tetramer (**Fig. 4C**). Furthermore, neither the kinetics nor the magnitude of either proximal (Syk, BTK) or distal (ERK1/2) signaling molecules was improved following exposure to GT8 60-mer, with the exception of BLNK phosphorylation (**Fig. 4D**). Collectively, these results demonstrate that when taking into consideration the stoichiometry of SOSIP trimer binding (n=3), and the cis-trans geometry of the binding sites in the streptavidin molecule (Fairhead et al., 2014) (used in tetrameric antigens) that likely allows divalent interaction on B cell surface, a minimum valency of n=2 is required for phospho-signaling and Ca-mobilization.

**Figure 4.**
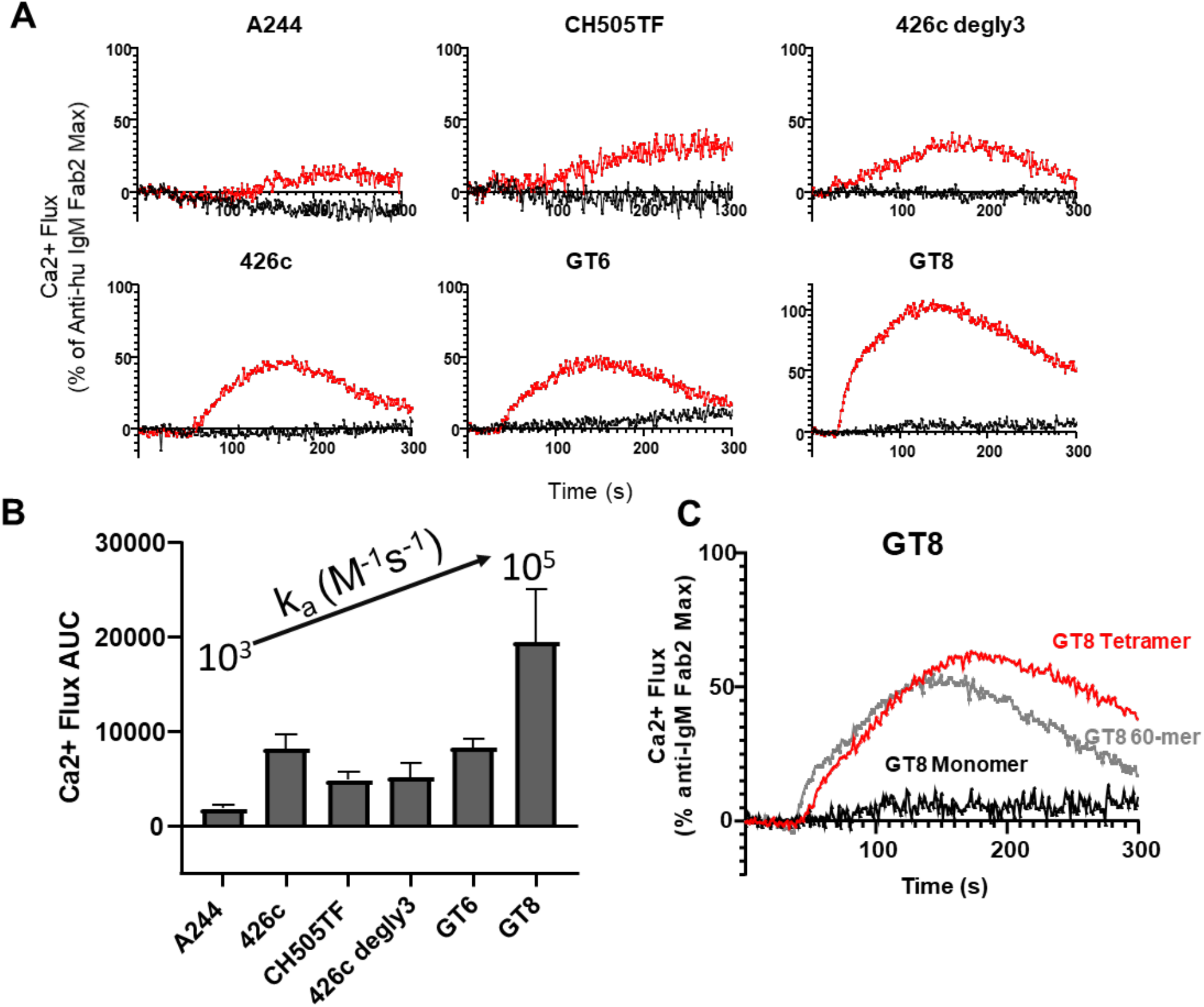
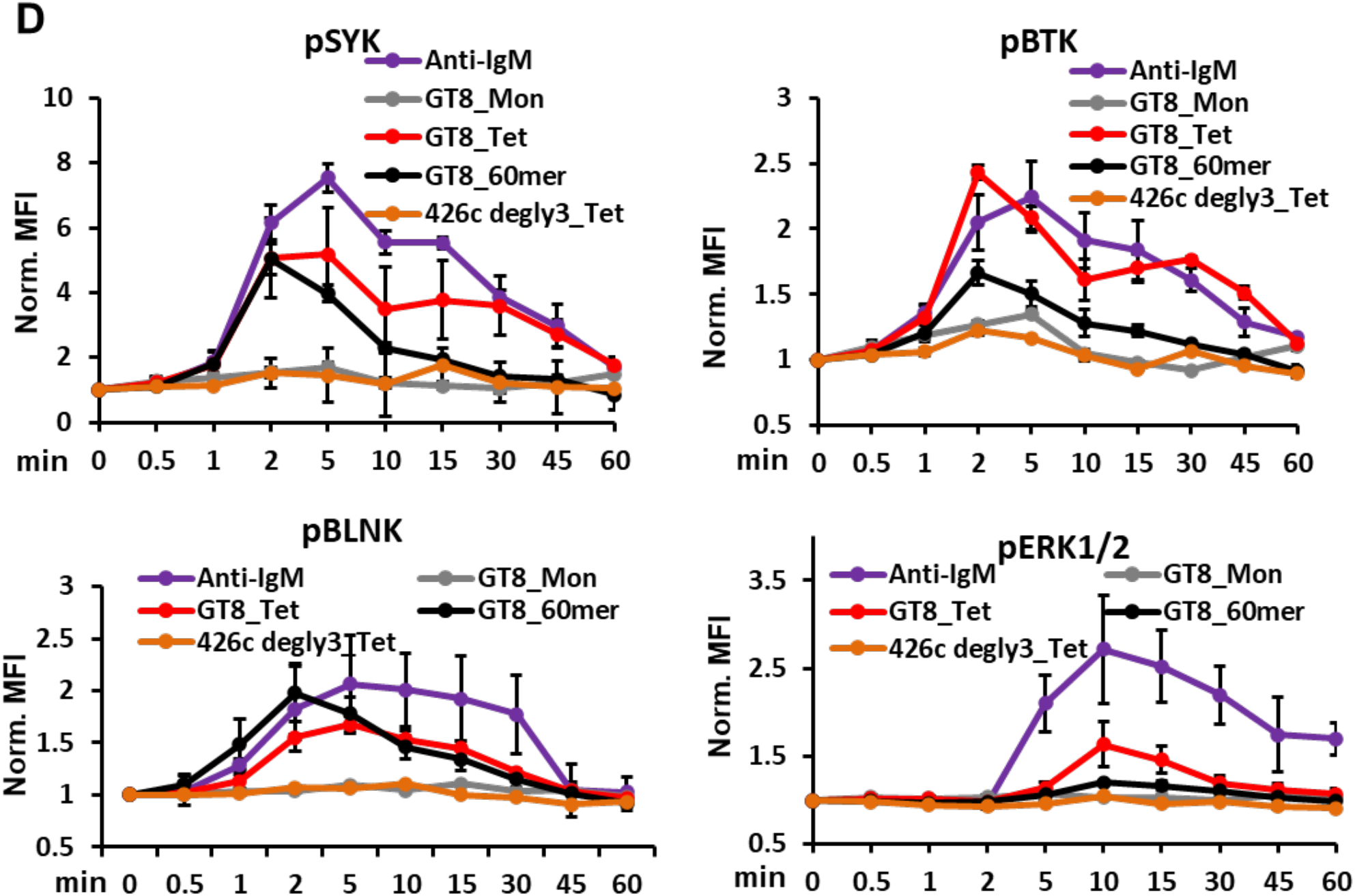
CH31 IgM B cell signaling by monomeric Env proteins. (**A**) Overlaid calcium flux responses of CH31 IgM Ramos cells mediated by a panel of monomeric antigens (black) and their multimerized (4-mer) counterparts (red). Monomeric proteins and their multimeric forms were prepared at the same per unit monomer concentrations for B cell activation (1μM). Results are presented as a % of the maximum anti-human IgM F(ab)2 induced responses and are representative of at least two measurements. (**B**) Bar graph of area under the curve (AUC) derived from CH31 IgM calcium flux responses mediated by multimerized proteins (4-mer). AUC values are an average of at least two measurements with error bars of the standard deviation and are plotted in order of slowest (left) to fastest (right) monomeric antigen association-rates. (**C**) Calcium flux responses of CH31 IgM Ramos cells induced by a high-order multimer (60-mer) of GT8 (gray) overlaid with the responses of the tetrameric (red) or monomeric (black) GT8 antigens. The 60-mer and 4-mer antigens were prepared at the same per unit monomer concentration of GT8 (250nM) and the concentration of GT8 monomer at 1μM. (**D**) Time-dependent phosphorylation of proximal and distal signaling molecules induced by monomeric Env proteins in different forms. The kinetics of proximal Syk, BLNK, Btk and the distal ERK1/2 phosphorylation in CH31 IgM Ramos cells were measured over 1h of exposure to either GT8 monomer (grey), GT8 tetramer (red), GT8 60-mer (black) and 426c degly3 (orange), with each protein at a concentration of 250 nM along with anti-human IgM (purple) at 30 nM. CH31 IgM cells were collected at each indicated time, fixed, permeabilized and then stained for intracellular level of phosphorylated Syk, BLNK, Btk or ERK1/2. Data plotted here are MFI values normalized to unstimulated sample MFI and are means and standard deviation of three separate experiments.

### Antigen binding half-life threshold for Ca-flux

The inability of monomeric Env proteins with varying binding k_a_ rates (10^3^ to 10^5^ M^-1^s^-1^) to trigger Ca-flux suggested that they dissociated too quickly, i.e., the antigen-bound BCR complex did not have long enough half-lives (t_1/2_) to activate B cells. The above consideration led us to investigate the impact of multimerization on the protein bound complex t_1/2_, as calculated from binding dissociation rates, k_d_ (**Fig. 5**). For monomeric proteins (GT8, GT6, 426c degly3, CH505TF) that bound with relatively faster association rates (>10^4^ M^-1^s^-1^), the t_1/2_ values of antigen-bound BCR complexes were less than 3 min, with the majority (3/4) of them binding with t_1/2_ less than 15 sec (**Figure 5C**). The above t_1/2_ values indicated that for B cell activation, there is a BCR-antigen half-life threshold that was not reached by any of these monomeric proteins. Tetramerization of each of the above monomeric proteins resulted in prolonged t_1/2_, ranging from about 5- to 100-fold (**Fig. 5A** **& 5C**), and resulted in the induction of Ca-flux by the tetramers (**Fig. 4A**). However, the magnitude of Ca-flux was not dependent on the t_1/2_ or affinity, but on binding association rate (**Fig. 4**). Thus, we observed no flux with A244 gp120 that bound with a slow association rate. A comparison of GT8 in tetrameric or 60-mer form showed that the higher valency multimeric interactions of the latter (with significantly prolonged half-lives i.e.∼hours) did not notably enhance either proximal or distal signaling over that observed with the GT8 tetramer (**Fig. 4D**). These results indicate that there is a half-life ceiling and the window of time from the threshold to the ceiling is a range spanning from less than a min (∼0.1 min) to less than an hour (∼44 min for GT8 tetramer), although a t_1/2_ of about 5 min (GT6 tetramer) was required for appreciable (50% of reference) flux (**Fig. 4**). The half-lives of the Env trimers (**Fig. 5C**) that gave Ca-flux without higher-order multimerization (**Fig. 2** **& S6**) provide further support to the above predicted ceiling of antigen-bound BCR complex half-lives.

**Figure 5.**
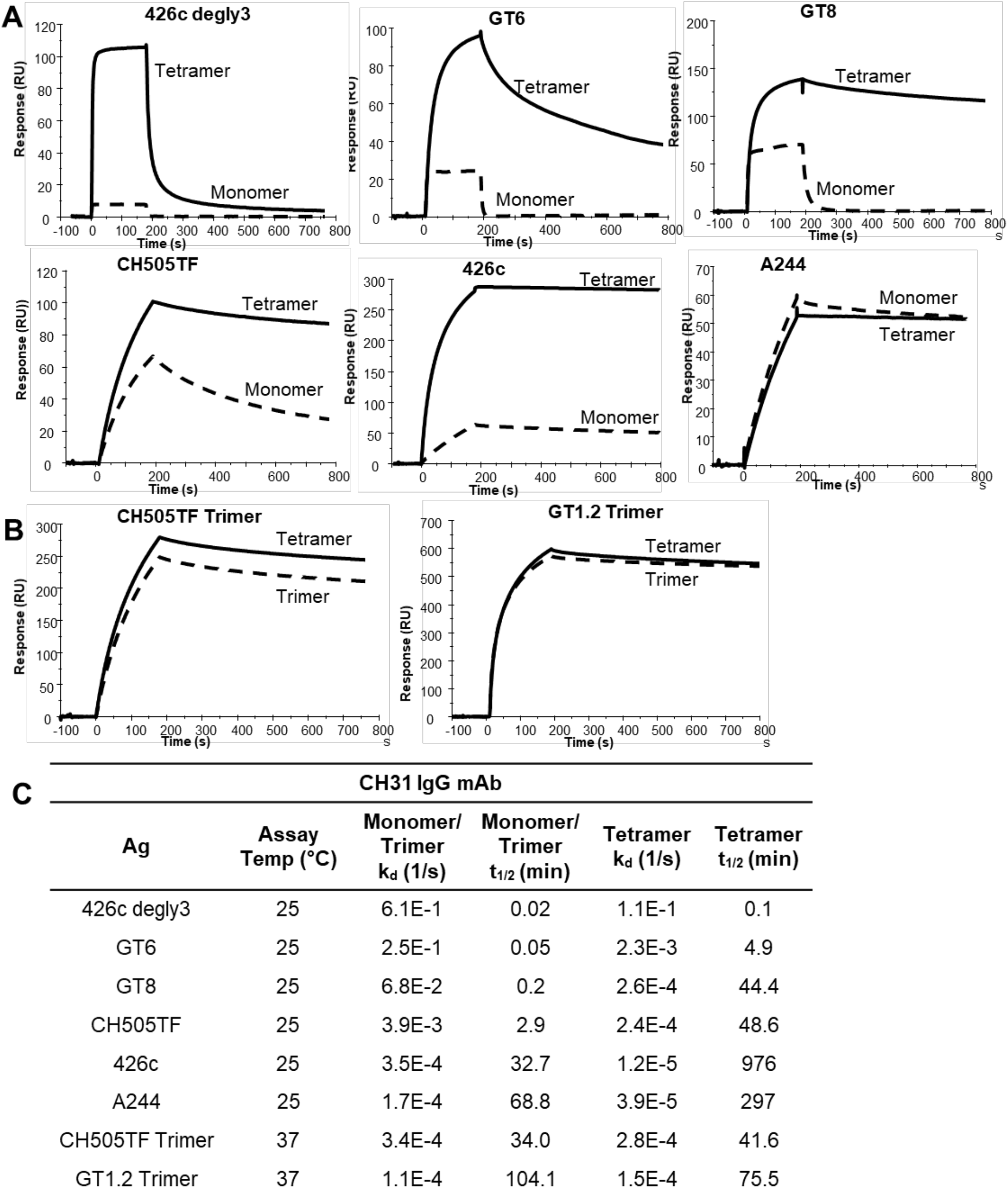
Effect of multivalent interactions on binding dissociation rates. (**A**) SPR Binding curves of monomeric proteins (dotted line) overlaid with the binding curves of multimerized (4-mer) forms (solid line) of the same proteins to CH31 IgG. Monomeric proteins and their multimeric forms were screened at the same per unit monomer concentrations for mAb binding. (**B**) SPR curves of trimeric antigens (dotted line) compared with the binding of the multimerized (4-mer) trimers (solid line). Trimeric proteins and their multimeric forms were screened at the same per unit trimer concentrations for mAb binding. (**C**) Env protein binding off-rates (k_d_, 1/s) and half-life (t_1/2_) of antibody-bound complex to CH31 IgG. Protein-antibody half-life (t_1/2_) were derived from k_d_ values using t_1/2 =_ ln(2)/k_d_. Binding curves as well as k_d_ and half-life measurements are representative of at least two measurements.

### Stronger BCR downmodulation induced by Env trimer with faster binding association rate

To investigate the role of Env binding kinetic rates on the endocytic activity of CH31 IgM BCR, we measured the ability of Env proteins to induce BCR downmodulation. We evaluated BCR downmodulation potential of two SOSIP trimers, CH505TF and GT1.2, that we demonstrated can induce Ca-flux in CH31 IgM cells. While similar BCR downmodulation was observed with both trimers at the highest concentration used (300 nM), dose response analysis showed that GT1.2 induced stronger BCR downmodulation with an EC_50_ value 7 times lower than CH505TF’s (**Fig. 6A**). Furthermore, GT1.2 showed an even larger difference in EC_50_ (18-fold) relative to CH505TF’s, when tested for BCR downmodulation in VRCO1 IgM expressing Ramos cells (**Fig. 6B**). Therefore, we conclude that for Env trimers with similar binding affinities, the strength of BCR downmodulation is dependent on the association kinetics, k_a_ of antigen binding.

**Figure 6.**
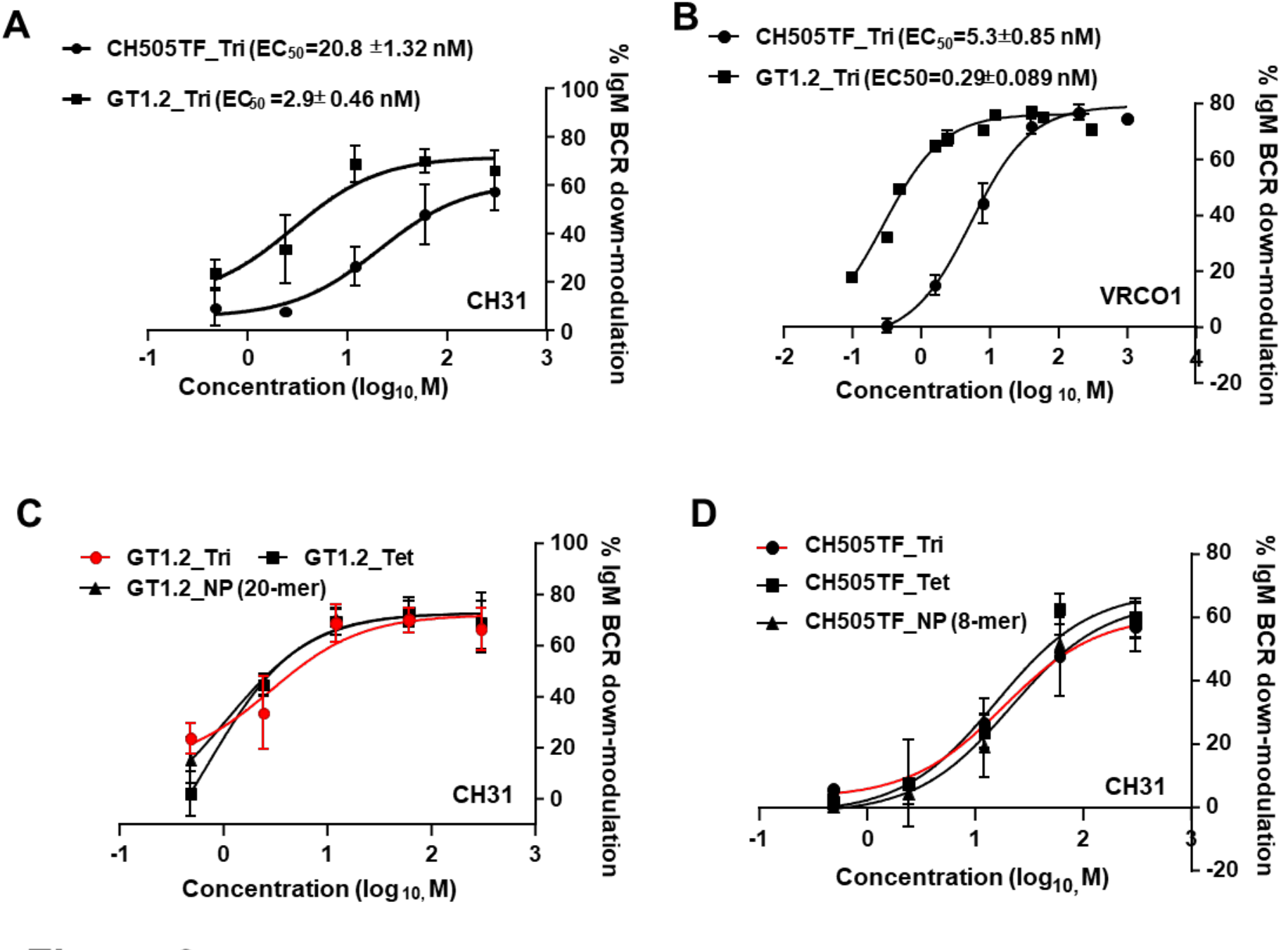
Antigen-binding induced BCR downmodulation. Dose response curves of CH505TF or GT1.2 trimer-induced %IgM BCR downmodulation following exposure of each protein to (**A**) CH31-IgM or (**B**) VRC01-IgM Ramos cells. Cells were stimulated with either Env trimers (CH505TF and GT1.2) over a concentration range of 0.48nM to 300nM or PBS for 1h at 37^0^C and subsequently stained for surface IgM BCRs and analyzed by flow cytometry. Percent of IgM BCR downmodulation for each concentration was obtained by the median fluorescence intensity (MFI) of the sample and PBS treated control (**C-D**) Dose-dependent % IgM BCR downmodulation of CH31 IgM BCRs following exposure to GT1.2 (**C**) or CH505TF (**D**), each protein in either trimer (circle), tetramer of trimers (rectangle) or higher-order multimeric nanoparticle (NP) forms (triangle, GT1.2 20-mer or CH505 8-mer). Data shows mean %IgM BCR downmodulation with standard deviation calculated from three independent experiments.

We next compared the impact of Env valency on BCR downmodulation in bnAb-expressing B-cells. When compared to the GT1.2 trimer, both GT1.2 multimeric forms (4mer and 20-mer) induced identical dose-dependent downmodulation profiles (**Fig. 6C**). Similarly, multimeric forms of the CH505TF SOSIPs mediated comparable BCR downmodulation relative to their standard trimeric forms (**Fig. 6D**). Thus, increasing the valency of SOSIP trimeric proteins had no enhancing or reducing effect on either early B cell activation events (Ca-mobilization) or BCR downmodulation.

### Antigen binding-induced internalization of monomeric and trimeric Env proteins

To investigate the role of binding kinetics on IgM BCR-mediated internalization of antigens, we measured the time course of antigen dissociation from the cell surface (surface antigen) and the level of internalized antigen (intracellular antigen) following exposure to Env proteins (**Fig. S9**). Exposure of CH31 IgM B-cells to GT6 tetramers (t_1/2_ = 4.9 min) resulted in a loss of surface antigen (∼80%) within 30 mins (**Fig. 7A**). The rapid decay of antigen from the cell surface was not associated with any detection of intracellular GT6 antigen (**Fig. 7A**), indicating that it was entirely due to the dissociation of surface BCR-bound complex without antigen internalization. CH31 IgM B-cell exposure to GT8 tetramers (t_1/2_ = 44.4 min) also resulted in loss of surface antigen within the first 30 min but required a longer time (90 min) to reach about 80% surface antigen loss (**Fig 7B**). However, unlike exposure with GT6, that with GT8 resulted in accumulation of intracellular antigen, beginning after 60 min, and further increasing at 120 min (**Fig. 7B**). Furthermore, while tetramers of SOSIP trimers CH505TF and GT1.2 both mediated surface antigen decay with kinetics comparable to that of GT8 tetramers (**Fig. 7C**), accumulation of intracellular antigen exposure was markedly pronounced in response to tetrameric GT1.2 relative to CH505TF. Thus, it is notable that the association rate difference between GT1.2 and CH505TF trimers was reflected in both B cell signaling and in BCR/antigen internalization. Together, these results indicate that an antigen must remain bound to BCR long enough to be internalized (t_1/2_ >5 min) but the degree of internalization is dependent on antigen binding association rate and not overall K_D_ value.

**Figure 7.**
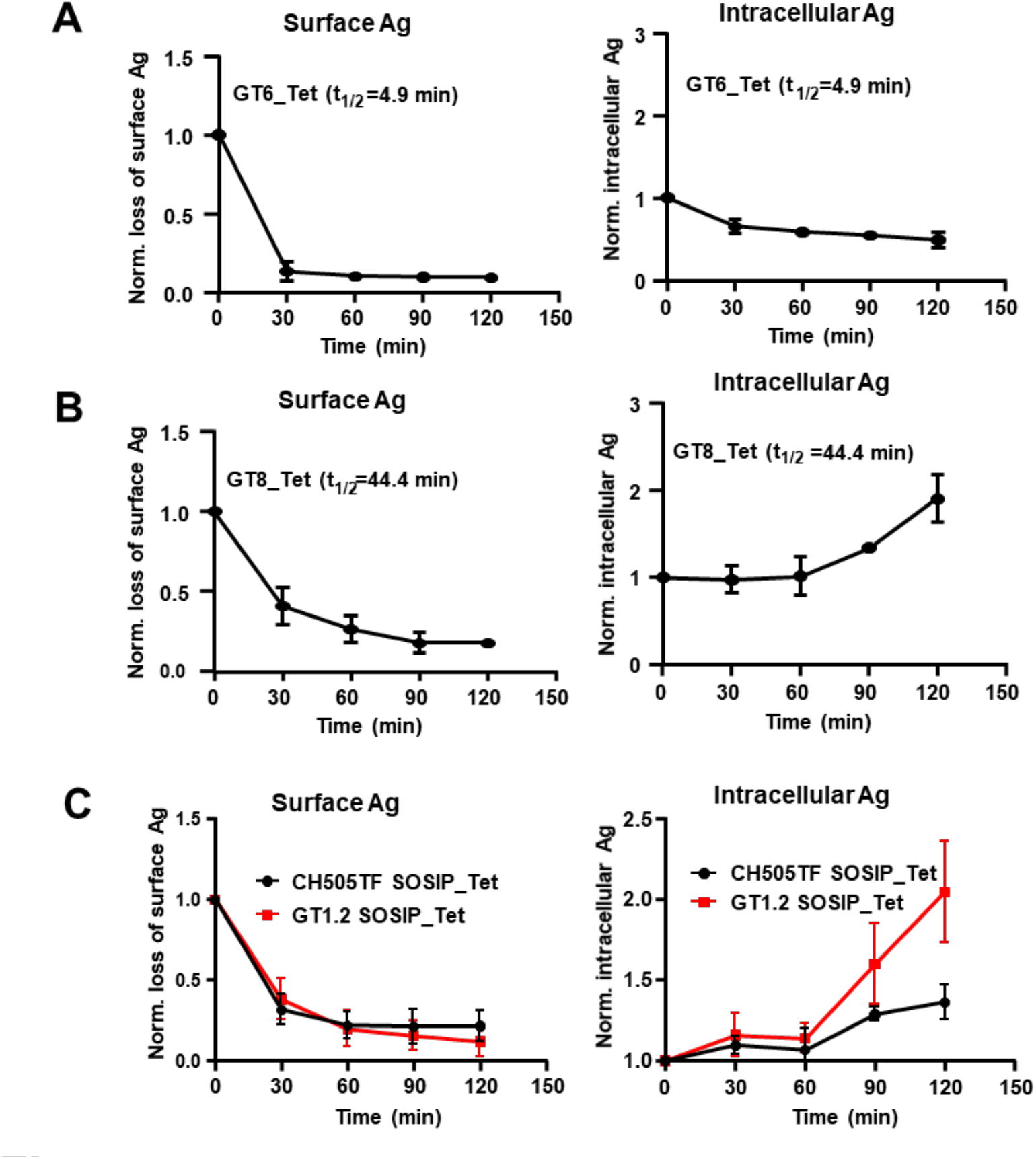
Dissociation of surface bound antigen and BCR-antigen internalization. The loss of CH31 IgM BCR bound antigen (left panel) and the level of intracellular Env proteins (right panel) following exposure to the indicated proteins over time. Surface loss and internalization of tetramers of GT6 **(A)**, GT8 **(B)** or (**C**) trimers (CH505TF, GT1.2) were measured by flow cytometric analysis following exposure of each protein to CH31 IgM Ramos cells over 2h period. Plotted data in the figure (**A-C**) are means of normalized MFI values of antigen treated cells to unstimulated cells or 0 min control calculated from three independent experiments. Half-life (t_1/2_) values of Env-CH31 bound complex are indicated in each plot. Data presented are means with standard deviation values from three independent measurements at each time-point.

## DISCUSSION

In this study, we report that B cells test antigen affinity by sensing the association rate and that the strength of activation increases with increasing association rate, regardless of the overall affinity (K_D_). In addition, our studies show that there is a half-life threshold for antigen-specific BCR activation and resulting BCR/antigen internalization, and while a minimal BCR-antigen dwell time is required for internalization, the magnitude of internalization is dependent on the antigen’s binding association rate. Thus, our studies define the optimal association rate range (k_a_ =10^4^ - 10^6^ M^-1^s^-1^), the minimal valency (n=2), and the threshold half-life required for B cell signaling (0.1 min) and BCR/antigen internalization (>5 min). These results support a kinetic rate dependent affinity discrimination for B cell activation and provides an explanation for the observed diversity in affinities as well as the lack of strict dependence of B cell responses to in-vitro measured antigen affinity.

The correlation between association rate and B cell activation described here using Ramos cells expressing CD4bs-directed bnAbs (CH31, VRC01) as BCRs was also observed in an *ex vivo* Ca-flux assay using primary naïve B cells from CH31 UCA knock-in mice (Verkoczy & Zhang, unpublished data). Furthermore, in an earlier study, the correlation between signaling and affinity was observed to be imperfect and led to the prediction that kinetic rates rather than equilibrium affinity might be the key binding parameter for B cell activation (Kouskoff et al., 1998). Using a large panel of Env proteins in different forms, we report here that the association rate is indeed the key predictor of cell signaling events leading to calcium mobilization. One notable finding in our study is that two Env trimers having similar, moderate binding affinities gave strikingly different responses both in B cell signaling and BCR/antigen internalization, while the high affinity trimer failed to induce a response. We further demonstrated that the two trimers inducing different B cell signaling kinetics bind with distinct thermodynamic mechanisms of interaction. In particular, as a measure of conformational change during binding, higher ΔCp values were found to be associated with slower k_a_ and weaker stimulation, both in the case of trimers as well as when comparing monomeric forms of the Env proteins (**Fig. S5**). Analogously, inclusion of ΔCp values to TCR binding half-life has been found to more accurately predict the strength of signaling in T cell responses (Krogsgaard et al., 2003). Our data indicates that sensing of antigen affinity by B cells has a qualitative and temporal basis and thus will require biophysical measurements of time-resolved association dynamics of antigen binding. In this regard, molecular dynamic simulation studies may be useful to model distinct interaction pathways and provide a better understanding of the association dynamics during antigen encounter and formation of the bound complex (Henderson et al., 2019).

An antigen may bind to a soluble form of antibodies differently than when it interacts with the membrane-bound IgM-BCRs (Iype et al., 2019). The differences we find in the kinetics and affinity of Ag-binding in the SPR analysis and the binding features to the IgM B cells suggest that the Ag binding of soluble monoclonal IgG antibodies and of the IgM-BCR complex on the B cell surface has quite different biophysical properties. When compared to the 3-dimensional (3-D) translational/rotational freedom of a soluble antibody, the IgM-BCR will be restricted to fewer degrees of freedom in the membrane-bound 2-D environment, and antigen binding may be dependent on the ‘local 2D-association rate’ (Tolar and Spillane, 2014). In addition, due to the oligomerization and presumably also a lateral shielding of the IgM-BCR on the B cell membrane its two Fab-arms may not be flexible, and the Ag binding site maybe stuck in a particular conformation that is only approachable by Ag with a high association rate. The interaction of high affinity Ag with the IgM-BCR, on the other hand, maybe dependent on an “induced fit” and alterations of the CDR loops that occurs only upon opening of the Fab arms. Thus, the interaction of high affinity Ag may require opening of the closed IgM-BCR oligomer in a time-dependent process. Indeed, it has been shown that the opening of the Fab arms can alter the conformation of the Ag binding site (Sela-Culang et al., 2012).

Using both trimeric and monomeric proteins, we also show that a minimal valency of n=2 is required for both activation and BCR/antigen internalization. The minimal valency for activation was also required for the GT8 monomer that bound with a half-life well above the minimal threshold (0.2 min), and is consistent with a recent report on the minimal valency needed to induce Ca-flux (Veneziano et al., 2020). Our study did not show any appreciable enhancement in signaling, in response to higher-order multimers (8-60 mers), either for monomeric or trimeric Env proteins. BCR responses may thus require “quantized multivalent interactions” (Vogelstein et al., 1982); however, dimeric or trimeric forms of the antigen may be sufficient for triggering BCR. The observed lack of enhanced triggering by higher-order multimers when compared to dimeric or trimeric forms provides support to the ‘Dissociation activation model” (Yang and Reth, 2010a, b) and is inconsistent with a strict requirement of clustering of monomeric BCRs through extensive multivalent interactions. Nevertheless, in the *in vivo* secondary lymphoid environment, higher-order nanoparticles or other particulate antigen forms could provide more efficient lymphatic trafficking, longer residency and enhanced MHC class II presentation, thereby improving immunogenicity.

Finally, our studies report on the role of BCR-antigen half-life in BCR/antigen internalization, a key step leading to antigen presentation and recruitment of T-cell help. While we observed that antigens bound to surface BCR long enough to be internalized, the requirement of the half-life threshold was found to be secondary to association rate. For monomeric antigens that bound with faster on-rates (k_a_> 1x10^4^ M^-1^s^-1^) but shorter half-life values (< 3 min), the antigen dwell time could be prolonged long enough for BCR/antigen through cis-divalent interactions following tetramerization to streptavidin. The converse, however, was not true, since an antigen with a slower binding dissociation rate (A244 gp120) and in tetrameric form failed to induce BCR/antigen internalization. Both A244 and 426c monomers bound with slow association rates (∼2x10^3^ M^-1^s^-1^) and with relatively long t_1/2_ (>30 min) (**Figure 5C**), but only 426c tetramers induced calcium mobilization (**Figure 4A**). An examination of the binding curves of monomeric and tetrameric forms of the two proteins showed that, unlike A244 tetramers, 426c tetramers showed greatly enhanced binding (∼4.5x more) when compared to their monomeric form (**Fig. 5A, S10**). This improved binding of 426c tetramers, despite the slow monomeric on-rate, represents an exception for a needed k_a_ threshold in our studied panel of antigens. However, while such enhanced multivalent interactions may be beneficial for activating B cells with weak affinity BCRs (presumably, by compensating for the low association-rates, via allowing them to make multiple productive contacts with BCR), it raises a concern that higher-order multimers could also provide enhanced competitiveness for triggering ‘off-target’ responses i.e. those to other B-cell epitopes likely to be found in more complex immunogens like Env proteins. Another caveat of our current study is that although we measured BCR/antigen internalization, presentation of the internalized protein derived peptides to MHC class II was not measured. Notably, the estimated BCR-antigen half-life range defining the threshold and ceiling (>5 – 34 min) for BCR/antigen internalization described here are similar to an earlier report on the role of k_d_ in antigen presentation to variant antigens that all bound with similarly fast k_a_ rates (Batista and Neuberger, 1998).

Our overall finding that B cells test affinity by sensing antigen binding association rate, has potentially important translational implications for vaccine research, and is especially important for diverse and antigenically heavily occluded pathogens like HIV. Induction of anti-HIV bnAb responses is highly challenging (Haynes and Mascola, 2017), and the parameters for BCR signaling and activation of bnAb precursor B cells remain to be well-defined. Our results suggest that designing Env proteins with improved association kinetics and not strictly based on equilibrium dissociation constants may have higher predictive value for ranking immunogen candidates in HIV vaccine studies that seek to stimulate and drive bnAb lineages. As one example of this, we have found in a separate study, that immunogens targeting a V3-glycan bnAb precursor (DH270 UCA) (Saunders et al., 2019) and designed to improve association kinetics, but not affinity, induced stronger Ca-mobilization in Ramos cells expressing BCRs of the bnAb UCA (Henderson, Azoitei, Alam et al., unpublished).

## Author Contributions

SMA conceived and designed the study and wrote the paper. MAH and KA co-wrote and edited the paper. MR and LV provided collaborative support, revised and edited paper. KA performed and analyzed SPR and calcium flux experiments, MAH phenotyped Ramos cell lines, performed and analyzed cell surface binding, phospho-signaling, and BCR/antigen downmodulation and internalization experiments. BW performed and analyzed thermodynamics experiments. KC prepared and purified Fab-trimer complexes and RJE performed and analyzed NSEM data. APK performed and analyzed antigenicity of Env proteins. JZ performed ex-vivo Ca-flux experiments. MR and YO assisted with the protocols for BCR/antigen downmodulation and internalization. DE developed cell lines, and MAH phenotyped, and sorted for high BCR expressing Ramos cell lines expressing CH31, VRC01 or VRC01 UCA-IgM BCRs. WR performed statistical analysis for correlation evaluation. All authors reviewed and edited the manuscript.

## METHODS

### Antigens

The following antigens were produced at the Duke Human Vaccine institute as previously described: eOD-GT6 (KX527852) (Jardine et al., 2013), eOD-GT8 (KX527855) (Jardine et al., 2013), 426c core WT (KX518323) (McGuire et al., 2016), 426c-degly3 core (KX518319) (McGuire et al., 2016), A244 D11gp120 (Alam et al., 2013), CH0505-Con D7 gp120 (Liao et al., 2013), CH505TFv4.1 SOSIP (Saunders et al., 2017), CH848 10.17DT SOSIP (Saunders et al., 2019), CH505TF SOSIPv4.1_SORTAv3_Ferritin (Jumaa et al., 2005), and CH848 DS SOSIP N133D_N138T_cSORTA_Ferritin (Paus et al., 2006). For biotinylation and subsequent tetramerization of these proteins, the envelope sequences were expressed with a C-terminal avidin tag (AviTag: GLNDIFEAQKIEWHE). eOD-GT8 d41m3 60mer (KX527857) (Jardine et al., 2013) was produced at The Scripps Research Institute. BG505 GT1.2 SOSIP and BG505 GT1.2 nanoparticle were produced at the University of Amsterdam and provided by Rogier W. Sanders. The production and purification of BG505 GT1.2 SOSIP was as described for previous soluble trimers (Pugach et al., 2015) while production for BG505 GT1.2 SOSIP nanoparticle was as described for previous nanoparticles (Brouwer and Sanders, 2019).

### Surface Plasmon Resonance (SPR) Affinity and Kinetic Rate Measurements

SPR derived kinetic rates (k_a_ and k_d_) and affinity measurements (K_d_) of monomeric proteins (germline-targeting (GT) monomeric forms of outer domain of gp120, 426c core proteins derived from the clade C 426c Env, and gp120 proteins) against CH31 IgG mAb were obtained using the Biacore S200 (Cytiva) instrument in HBS-EP+ 1X running buffer. A Protein A chip (Cytiva) was used to capture CH31 IgG mAb to a level of 200-300RU on flow cells 2, 3, and 4 for all proteins, except for 426c degly3 where the capture was approximately 700RU. A negative control antibody, Ab82, was captured onto flow cell 1 to approximately 300-400RU for reference subtraction. Proteins were injected over the sensor chip surface using the single cycle injection type. Six sequential injections of the samples diluted from 25nM to 6000nM were injected over the captured CH31 IgG mAb at a flow rate of 50uL/min for 120s per injection. The single cycle injections were followed by a 600s dissociation and regeneration with a 20s pulse of Glycine pH1.5. The Biacore S200 Evaluation Software (Cytiva) was used to analyze results. Binding to the negative control antibody, Ab82, as well as buffer binding were used for double reference subtraction and accounted for both non-specific binding and signal drift. Curve fitting analyses were performed using the 1:1 Langmuir model with a local Rmax for all antigens except CH505TF gp120. CH505TF gp120 titration curve was fit using the heterogeneous ligand model with faster kinetic rate parameters reported. The reported kinetic rates and affinities are representative of 3 data sets.

SPR affinity measurements (K_d_) and kinetic rates (k_a_ and k_d_) for the trimeric gp140 proteins were obtained using the Biacore S200 or T200 instrument (Cytiva) in HBS-EP+ 1X running buffer. Biotinylated SOSIP trimer proteins were immobilized to a level of 100-300RU onto a SA (streptavidin) sensor chip (Cytiva). CH31 IgG Fab was used as the analyte. CH31 Fab was diluted from 50nM to 1500nM and injected over the sensor chip surface using the single cycle injection type. Five or six sequential injections of the CH31 Fab were injected at a flow rate of 50uL/min for 120s per injection and were followed by a 600s dissociation period. Regeneration with a short pulse of Glycine pH2.0 was used for all trimers. Binding to a blank streptavidin surface as well as buffer binding were used for double reference subtraction, non-specific binding, and signal drift. A 1:1 Langmuir model with a local Rmax was used for the CH31 Fab curve fitting analysis against CH848 and CH505TF trimers. The heterogeneous ligand model was used for CH31 Fab binding to GT1.2 trimer with the faster kinetic rates reported. The reported kinetic rate and affinity values are representative of 2 data sets.

### Antigen Biotinylation and Tetramerization

Monomeric and trimeric proteins were biotinylated and tetramerized as previously described (Bonsignori et al., 2018). Biotinylation was performed using a BirA biotin-protein ligation kit (Avidity) on proteins produced with a C-terminal avidin tag sequence. Following addition of enzyme kit reagents, proteins were agitated at 900 rpm for 5h at 30°C. Excess biotin was removed using 10kDa MW spin columns (Amicon) for monomeric proteins and 100kDA MW spin columns (Amicon) for trimeric proteins. Proteins were transferred to the spin columns after incubation and five washes were performed in PBS 1X pH7.4 (Gibco). Tetramerization of monomeric and trimeric proteins was accomplished with streptavidin (Invitrogen). A 4:1 molar ratio of protein to streptavidin was used and to maximize streptavidin site occupancy; streptavidin was added in a stepwise manner to the protein. The appropriate volume of streptavidin was added 5x every 15 min followed by agitation at 900rpm and 23°C. The final molarity of the protein was calculated based on number of moles used in the reaction as well as the total volume of protein and streptavidin combined.

### Calcium Flux Measurements with Ramos cells

Calcium flux experiments were performed using the FlexStation 3 Microplate Reader (Molecular Devices) and Softmax Pro v7 software (Molecular Devices) in conjunction with the FLIPR Calcium 6 dye kit (Molecular Devices) as previously described (Dal Porto et al., 2002). On the day of the experiments, a cell count for the Ramos cells was performed with a Guava Muse Cell Analyzer (Luminex) to ensure cell viability was greater than 95% and to calculate the volume of cells needed for a concentration of 1x10^6^cells/mL. The appropriate volume of cells was then pelleted at 1500rpm for 5 minutes after which the supernatant was decanted and the cells were resuspended at a 2:1 ratio of RPMI media (Gibco) + FLIPR Calcium 6 dye (Molecular Devices). The cells were plated in a clear, U-bottom 96-well tissue culture plate (Costar) and incubated at 37°, 5% CO_2_ for 2h. Antigens were separately diluted down to a concentration of 2uM in 50uL of the 2:1 ratio of RPMI media (Gibco) + FLIPR Calcium 6 dye (Molecular Devices) and plated in a black, clear bottom 96-well plate. The final concentration of antigen would be 1uM based on the additional 50uL of cells added during the assay. A positive control stimulant, Anti-human IgM F(ab’)_2_ (Jackson ImmunoResearch) was also included in the antigen plate. Using the FlexStation 3 multi-mode microplate reader (Molecular Devices), 50uL of the cells were added to 50uL of protein or Anti-human IgM F(ab’)_2_ diluted in RPMI/dye and continuously read for 5min. Calcium flux results were analyzed using Microsoft Excel and GraphPad Prism v9. The relative fluorescence of a blank well containing only the RPMI/dye mixture was used for background subtraction. Once subtracted, the antigen fluorescence was then normalized with respect to the maximum signal of the IgM control and calcium flux values were presented as a percentage (% of Anti-hu IgM Fab2 Max). Calcium flux data are representative of at least 2 measurements for the CH31 Ramos cell line and one measurement for the VRC01 and VRC01 UCA Ramos cell lines.

### Surface Plasmon Resonance of Tetrameric Antigens

SPR binding curves of monomeric proteins compared with their tetrameric counterparts were obtained using the Biacore S200 instrument (Cytiva) in HBS-EP+ 1X running buffer. A Protein A chip was used to capture CH31 IgG mAb to a level of 220-320RU on flow cells 2, 3, and 4. A negative control antibody, Ab82, was captured on flow cell 1 to 300-400RU and was used for reference subtraction. Monomeric proteins and their tetrameric counterparts were diluted down to equivalent concentrations of 1000nM per unit of monomer, except for GT6 and 426c degly3 which were diluted down to 2000nM. These samples were injected at an assay temperature of 25°C over the antibody captured sensor surfaces for 180s at 30uL/min using the high-performance injection type. The trimeric CH505TF and GT1.2 proteins as well as their tetrameric versions were diluted down to 250nM per unit trimer and then injected over the antibody captured surface at 37°C for 180s at 30ul/min. For all antigens, a dissociation period of 600s followed the sample injection and the Protein A surface was regenerated with a 20s pulse of Gly pH1.5. After reference surface and buffer binding subtraction, curve fitting analyses (BIAevaluation Software) were performed on the single dose curves to measure the concentration independent dissociation rate (k_d_, 1/Ms), which was then used to measure the half-life (t_1/2_) of each antigen-antibody interaction. The half-life was calculated using the following equation: t_1/2 =_ ln(2)/k_d_. The reported k_d_ and t_1/2_ values reported are representative of 3 measurements.

### BCR downmodulation in CH31 IgM Ramos

For analysis of antigen induced downmodulation of CH31 IgM BCRs, Ramos cells were treated with a panel of HIV-1 env proteins in a concentration gradient manner. Approximately 2.5x10^5^ cells were challenged for each treatment and changes in surface BCRs expression were measured after 1hr at 37°C. Control samples were PBS treated and incubated for 1 hr at 37°C. Immediately after incubation, cells were placed on ice to stop the reaction and washed using cold PBS/1%BSA. For the detection of surface IgM, cells were incubated with anti-IgM-Fab-FITC for 30 mins in cold. Unbound antibodies were removed after washing with PBS/1%BSA, resuspended in 2% formaldehyde in PBS and analyzed by BD LSRII flow cytometer. Data from 10,000 single events were exported as FCS-3.0 format, analyzed with FlowJo software (version 10.8.1). Percent of IgM BCR downmodulation was calculated by the median fluorescent intensity (MFI) of the treated sample and control using the formula: % IgM BCR downmodulation= 100-((MFI of treated sample/MFI of PBS control) X100. Data plotted in the graph represents mean %IgM BCR downmodulation with standard deviation at any given protein concentration from triplicate experiments.

### Antigen internalization assay in CH31 IgM Ramos

For the detection of tetrameric antigen internalization by CH31 IgM BCRs, 5x10^6^ Ramos cells were incubated with Env proteins in PBS on ice for 20 mins. Cells were then washed by PBS, resuspended in 2 ml PBS/2% FCS and transferred to 37°C. Approximately, 100 ul of cells were harvested at each time points (0, 30, 60, 90, 120 mins) and placed in four separate flow tube (containing 4% formaldehyde) for the detection of surface IgM, surface proteins, blocked proteins and intracellular protein respectively. Cells were placed on ice for 10 mins and 30 mins at room temperature to fix. Fixed cells were washed by PBS/1%BSA and incubated with anti-IgM-Fab-FITC, anti-SA-PE and anti-SA for the detection of surface IgM, SA-conjugated tetrameric Env proteins, and blocking of surface SA-conjugates, respectively. Efficiency of surface blocking of SA-binding sites were confirmed after incubation with anti-SA-PE. For detection of intracellular SA-conjugated tetrameric proteins, surface blocked cells on fourth flow tubes were fixed 30 mins in IC Fixation buffer (#00-8222-49, Invitrogen), permeabilized after washing twice by permeabilization buffer (Invitrogen) and then incubated with anti-SA-PE to stain intracellular Env proteins. After 30 mins incubation at RT, cells were washed twice by permeabilization buffer to remove unbound antibodies. All cells after antibody incubation were washed and resuspended in PBS/1%BSA (0.05% azide) buffer and analyzed in BD LSRII Flow cytometer. The loss of surface Env proteins and intracellular Env protein tetramer over time were measured after normalizing MFI value of any given timepoints by the unstimulated or 0 min sample MFI value (considered no internalization).

### Antigen binding analysis in CH31 IgM Ramos cells

Time course binding of Env antigens to IgM BCRs were analyzed using flow cytometer. For that 2.5x10^5^ cells were incubated with HIV-1 antigens in PBS/2% FCS for the indicated period of times on a 37°C thermomixer (Eppendorf). Cells were then fixed with IC fixation buffer on ice for 30 mins and at RT for additional 30 mins. Fixed cells were incubated with fluorophore conjugated anti-SA and anti-biotin antibody to detect SA-conjugated tetrameric antigens and biotinylated antigens respectively. After incubation cells were washed and resuspended in PBS/1%BSA (0.05% azide) buffer and analyzed in BD LSRII Flow cytometer. The normalized value of MFI for each protein at a given time points calculated using PBS treated sample as negative control.

### Phospho-flow kinase analysis in CH31 IgM Ramos

Phosphorylation of three proximal kinases/adapter proteins (pSyk, pBtk, pBLNK) and one distal kinase (pERK1/2) were analyzed in CH31 IgM Ramos cells for an hour stimulation by HIV-1 Env antigens. A total of 2.5x10^6^ cells were washed in PBS, incubated with each antigen at certain concentration in PBS/2% FCS and transfer to a 37°C thermomixer (Eppendorf). Cells were collected after each period of incubation and fixed in IC fixation buffer on ice for 30 mins and at RT for additional 30 mins. Fixed cells were rinsed twice with permeabilization buffer (Invitrogen) and incubated with PE-Cy7 anti-Zap 70 phospho (Tyr 319)/Syk phosphor (Tyr 352) (#683708, clone: 1503310), BV421 mouse anti-human Btk (pY223)/Itk (pY180) (#564848, clone: N35-86, BD Biosciences), Alexa Fluor 488 mouse anti-human BLNK (pY84) (#558444, clone: J117-1278, BD Biosciences), and APC anti-ERK1/2 phospho (Thr202/Tyr204) (#369522, clone: 6B8B69) at RT for 1 hour. Cells were then washed twice by permeabilization buffer to remove excess antibodies and resuspended in PBS/1%BSA (0.05% NaN3) buffer. Data were collected for 10,000 single events using BD LSRII flow cytometer and analyzed using FlowJo software (v.10.8.1). Normalized MFI for each timepoints were obtained dividing treated sample MFI by unstimulated or 0 min sample MFI.

### Statistical Methods

To assess potential correlations, Kendall’s Tau analysis was used due to the small sample size. The alpha level was set at 0.05 and there were no adjustments made for multiple correlation evaluations. The analysis was conducted using SAS 9.4 software.

## Supplemental Information

Supporting Figures (S1-S10) are provided in the Supplemental Information document.

## Acknowledgments

We thank Dr. Bart Haynes, DHVI, Director of CHAVD (Duke Consortia) for providing facility resources, scientific advice and critical comments. We are grateful to Rogier Sanders, Ronald Derking (Amsterdam UMC) for providing GT1.2 gp140 trimer and NPs, William Schief, Bettina Groschell (Scripps Research) for GT Env proteins (monomers and multimers). We thank Kevin Saunders, Elizabeth Donahue (DHVI) for expressing and purifying Env gp120 and gp140 trimers and NPs. We thank the DHVI teams from BIAcore Facility, Flow cytometry, Protein Expression (Kevin Saunders) Center and the DHVI Finance and administrative teams for their support.

Research reported in this publication was supported by the National Institute of Allergy and Infectious Diseases (NIAID) of the National Institutes of Health (NIH) under Award Number R01AI145656 (PI: SMA). The content is solely the responsibility of the authors and does not necessarily represent the official views of the National Institutes of Health

---

**Figure S1.**
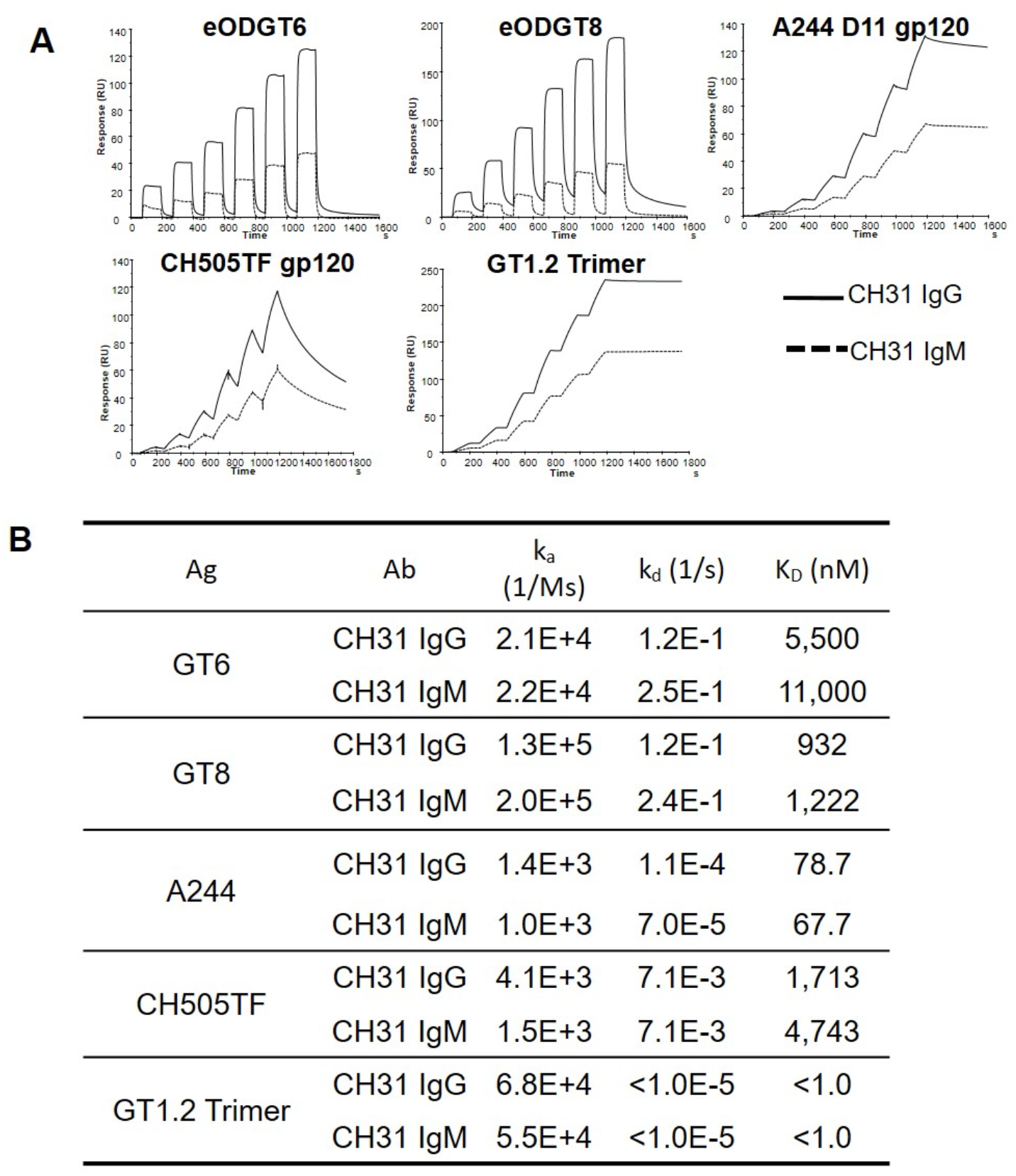
Affinity and kinetic rates of Env proteins binding to CH31 IgM mAb compared to CH31 IgG. (**A**) SPR single cycle kinetic binding profiles of monomeric (or trimeric) antigens against CH31 IgG and CH31 IgM. Plots are representative of two measurements. (**B**) Summary of the overall binding affinities (K_d_, nM) as well as the individual kinetic rate measurements (k_a_ and k_d_) of each antigen with either CH31 IgG or CH31 IgM. Values are representative of two data sets. A comparison of the binding results of the monomeric antigens (GT6, GT8, A244, and CH505TF) as well as the trimeric antigen, GT1.2, suggested that the affinities and kinetic rate parameters were similar for both isoforms of CH31. **Methods:** SPR derived affinities and kinetic rates of the antigens against CH31 IgG and CH31 IgM were obtained using the Biacore S200 (Cytiva) instrument in HBS-EP+ 1X running buffer. CH31 IgG and CH31 IgM were directly immobilized onto two flow cells of a CM5 chip to a level of approximately 2000RU while a negative control antibody, Ab82, was immobilized onto flow cell 1 to a similar response level for reference subtraction. Six sequential injections of each antigen diluted from 5nM to 10,000nM were injected over the immobilized IgG and IgM mAbs at a flow rate of 50uL/min for 120s per injection. The single cycle injections were followed by a 600s dissociation and regeneration with a 20s pulse of Glycine pH2.0. The Biacore S200 Evaluation Software (Cytiva) was used to analyze results. Binding to the negative control antibody, Ab82, as well as buffer binding were used for double reference subtraction and accounted for both non-specific binding and signal drift. Curve fitting analyses were performed using the 1:1 Langmuir model with a local Rmax for all antigens except CH505TF. The CH505TF gp120 titration curve was fit using the heterogeneous ligand model with faster kinetic rate parameters reported. The reported kinetic rates and affinities are representative of 2 data sets.

**Figure S2.**
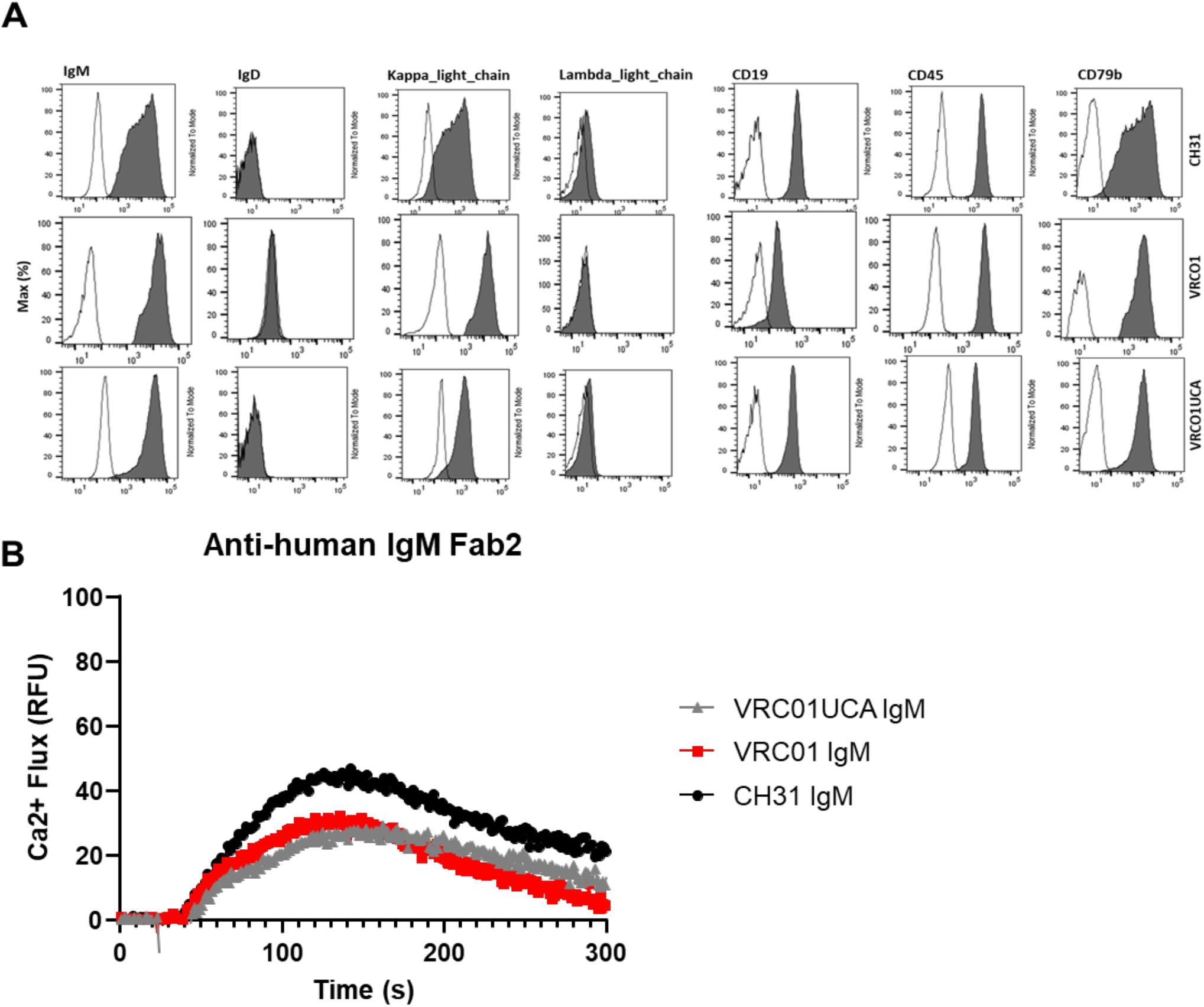
Ramos cell line phenotypic data and activation with control stimulant. (**A**) Representative flow cytometry plots showing expression of surface markers in bnAbs IgM (CH31, VRCO1 and VRCO1UCA) expressing ramos cells (dark) with unstained control (clear). Around 1.0x10^6^ cells were fixed and stained for the surface expression of IgM, IgD, k-light chain, ʎ-light chain, CD19, CD45, CD79b using fluor conjugated antibody and analyzed by BD LSRII flow cytometer. (**B**) Calcium flux of the following 3 Ramos cells lines: VRC01UCA IgM (gray), VRC01 IgM (red), and CH31 IgM (black) by the positive control stimulant Anti-human IgM F(ab)’2 at 50ug/mL. The fluorescence of a blank well containing only RPMI/dye was used for background subtraction. The calcium flux profiles are representative of at least two measurements and demonstrate the functionality of the three cell lines. **Methods:** Calcium flux experiments were performed as described in the methods section under “Calcium Flux Measurements with Ramos cells”.

**Figure S3.**
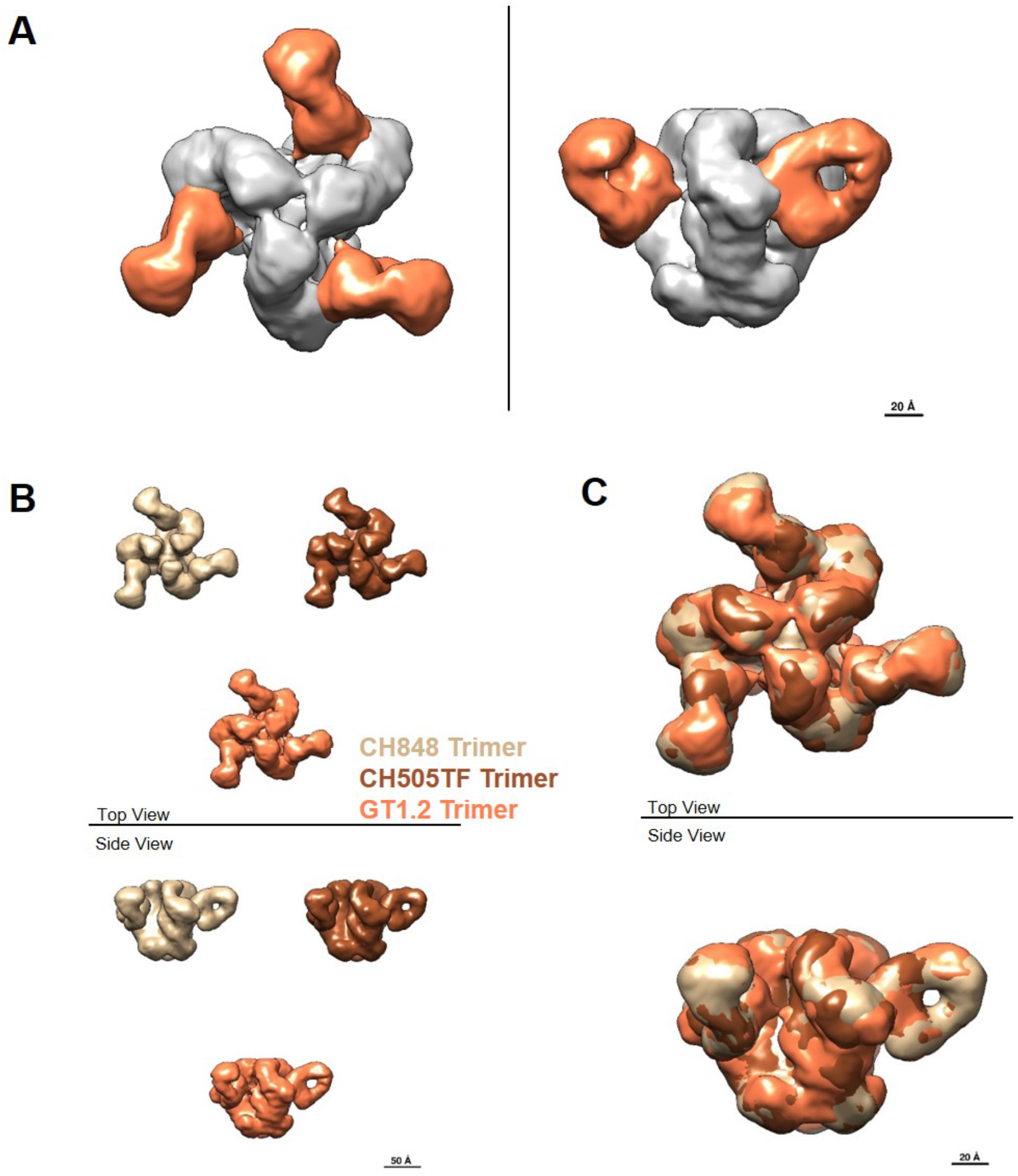
Negative stain-EM structures of CH31 Fab complexed to three different SOSIP trimers. (**A**) Structure of CH31 Fab bound to GT1.2 trimer, shown in top view and side view, with the density corresponding to the Env trimer colored gray and the density corresponding to the three Fabs colored orange. (**B**) Side by side comparison of the structures of CH31 Fab bound to CH848, CH505TF and GT1.2 trimers, colored tan, brown and orange, respectively. (**C**) Superposition of the three maps shown in **B** shows that at this resolution (∼20 Å) there is no significant difference in the observed epitope or binding angle of CH31 to the three different SOSIP trimers. **Methods:** **Fab-SOSIP Complex Preparation and Purification** Complex production was achieved by adding 15x Molar excess monoclonal Fab to 20ug trimer. This was then incubated overnight at room temperature. Fab-trimer complex was fixed by addition of glutaraldehyde to a final concentration of 5mM and then incubated for 10min at room temperature. Excess glutaraldehyde was quenched by adding sufficient 1M Tris and incubated for 10min at room temperature. Complex was purified through SEC by loading on a Superose 6 increase 10/300 column using a 100 μL loop, and run at 0.5 mL/min using an Äkta Pure system (GE Healthcare). Fractions were collected and complex peak pooled and concentrated by 10kDA cutoff centrifugal filtration (Amicon/EMD Millipore). Fab-SOSIP complex formation and quality was further accessed by SDS PAGE gel electrophoresis, where ∼1 μg complex/lane was loaded on a 4%–15% TGX Stain free gel (BioRad) in both reducing and non-reducing conditions, and run at 200 V in Tris/Glycine/SDS buffer. Bands were visualized using Gel Doc EZ imager (BioRad), and the size of the fragments assessed by a protein standard ladder (BioRad). **Negative stain electron microscopy** Purified sample of each Fab-trimer complex was diluted to 100 µg/ml with HEPES-buffered saline (20 mM HEPES, 150 mM NaCl, pH 7.4) augmented with 5% glycerol and applied to a glow-discharged carbon-coated EM grid for 8-10 second. Sample was then blotted and stained with 2 g/dL uranyl formate for 1 min, blotted and air-dried. Grids were examined on a Philips EM420 electron microscope operating at 120 kV and nominal magnification of 49,000x, and 100-130 images were collected on a 76 Mpix CCD camera at 2.4 Å/pixel. Images were analyzed by 2D class averages and 3D reconstructions calculated using standard protocols with Relion 3.0 (Zivanov et al. 2018. *eLife*. 7:e42166).

**Figure S4.**
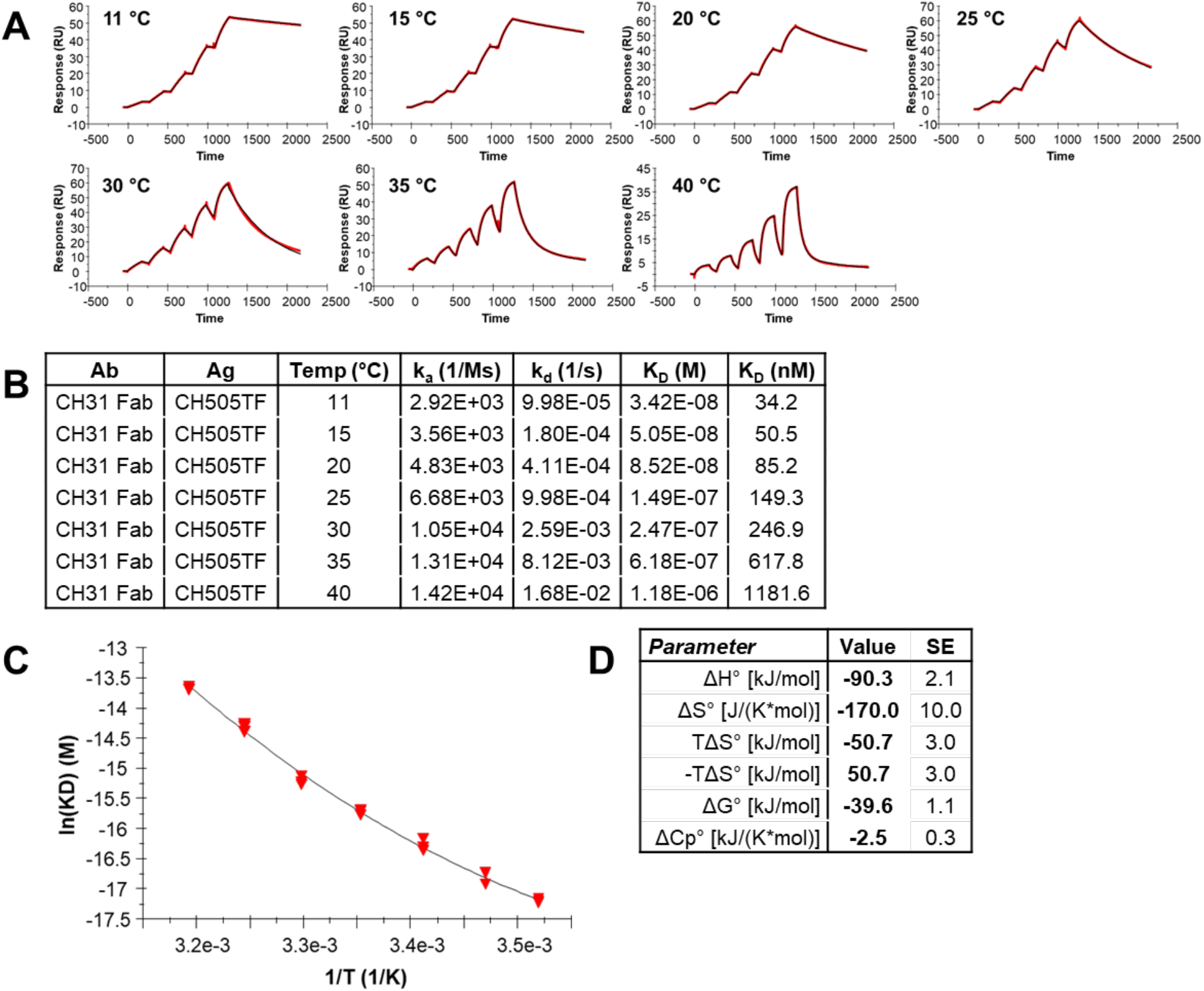
Thermodynamic measurement of CH31 Fab binding to the CH505TF trimer by surface plasmon resonance (SPR). (**A**) Single-cycle kinetic titration sensorgrams of CH31 Fab binding to the CH505TF SOSIP trimer recorded at varying temperatures. Plots are representative examples of 3 replicate measurements. (**B**) Table of association rate constants (k_a_), dissociation rate constants (k_d_), and affinity constants (K_D_) recorded at varying temperatures. Values are reported as the average of 3 replicate measurements. (**C**) Plot of the natural logarithm of affinity (K_D_) versus the reciprocal temperature and non-linear van’t Hoff fit. (**D**) Table of thermodynamic parameters obtained from non-linear van’t Hoff analysis. Values are reported as the average and standard deviation of 3 replicate measurements. **Methods**: Thermodynamic analyses of CH31 binding to trimeric and monomeric antigens were performed by surface plasmon resonance (SPR) on a Biacore T200 platform (Cytiva Life Sciences) at varying temperatures in HBS-EP+ running buffer (10 mM HEPES, 150 mM NaCl, 3 mM EDTA, 0.05% Surfactant P20, pH 7.4). For trimeric antigens, biotinylated SOSIPs (50 µg/mL) were captured on streptavidin sensor surfaces at 5 µL/min to densities of 230-420 RU. Affinity measurements were performed by sequential titration of 5 concentrations of CH31 Fab [62.5-1000 nM (GT1.2), 100-1600 nM (CH505TF), 300-2400 nM (CH848)] over the SOSIP surfaces for 150-180 seconds per injection followed by a dissociation period of 900-2700 seconds at 30-50 µL/min. Surfaces were regenerated between Fab titrations with 10 mM glycine-HCl pH 2.0 for 40 seconds at 30-50 µL/min. Titrations were repeated for each trimer at a minimum of 7 different temperatures ranging from 11-45 °C. For monomeric antigens, CH31 IgG mAb was captured on Protein A sensor surfaces at 5 µL/min to densities of 120-670 RU. Affinity measurements were performed by sequential titration of 5 concentrations of GT6 (625-10000 nM), GT8 (62.5-750 nM), or A244 (250-4000 nM) over a CH31 IgG surface for 180 seconds followed by a dissociation period of 720-1500 seconds at 50 µL/min. Surfaces were regenerated between antigen titrations with 10 mM glycine-HCl pH 1.5 for 45 seconds at 50 µL/min. Titrations were repeated for each antigen at 6 different temperatures ranging from 10-40 °C. Affinities were calculated using a steady state affinity model, a 1:1 kinetics model, or the fast components of a heterogeneous ligand kinetic model. Non-linear van’t Hoff analyses were used to derive binding enthalpy (ΔH), entropy (ΔS), free energy (ΔG), and heat capacity (ΔC_p_).

**Figure S5.**
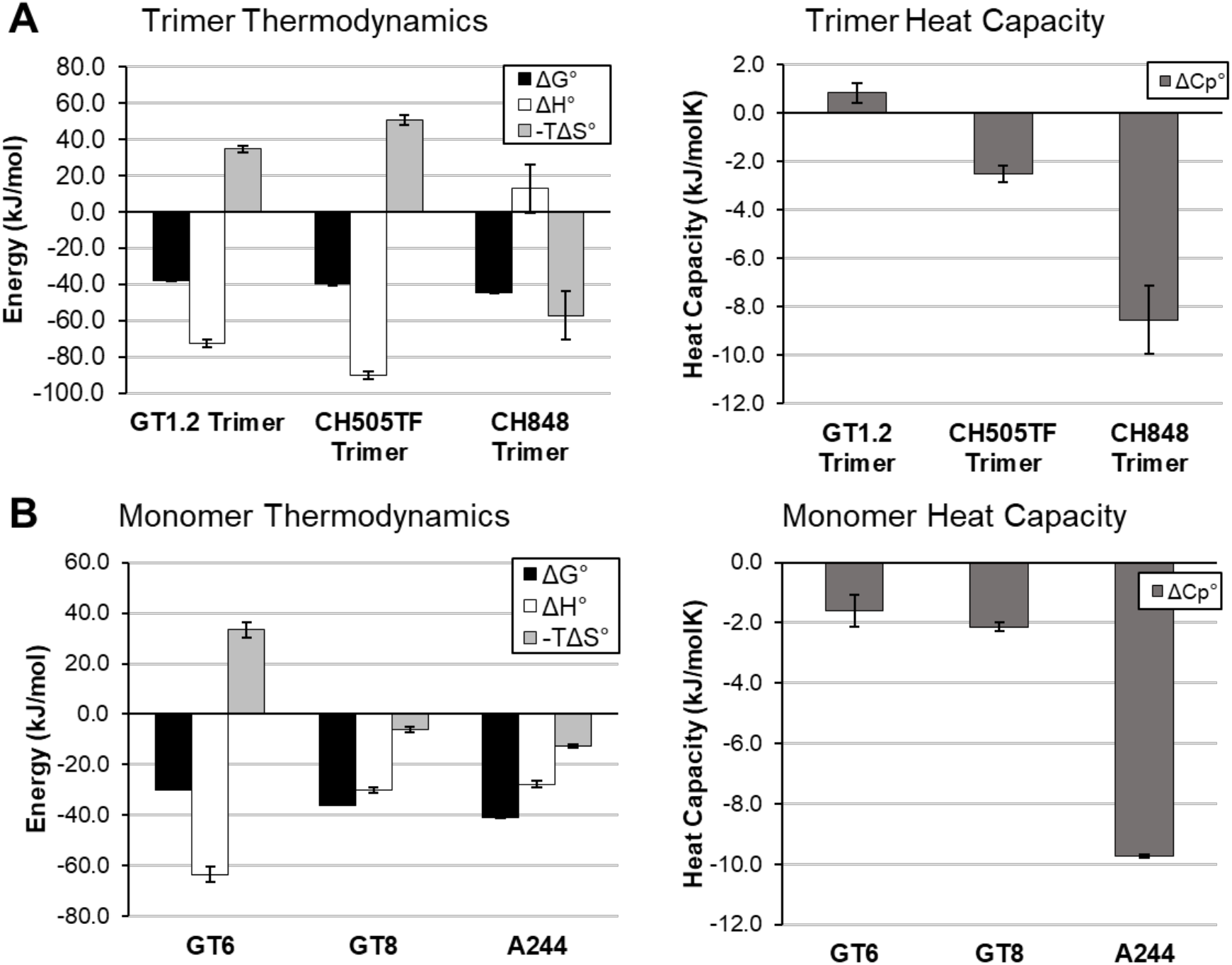
Thermodynamics of CH31 binding to SOSIP trimers and monomeric antigens. (**A**) Thermodynamic analysis of CH31 Fab binding to GT1.2, CH505TF, and CH848 trimers revealed markedly different binding profiles. The similar affinity GT1.2 and CH505TF trimers exhibited entropy-enthalpy compensation where the interaction was driven by strongly favorable binding enthalpies (ΔH < 0) and offset by unfavorable binding entropies (-TΔS > 0). The faster on-rate (k_a_ = 2.2x10^5^ M^-1^s^-1^) GT1.2 trimer displayed a relatively lower entropic hurdle (-TΔS) and a negligible ΔC_p_, which suggests that minimal conformational changes occur during the association process. Therefore, the GT1.2 trimer and CH31 Fab exist in unliganded states that require no conformational rearrangement to form a productive encounter complex. Alternatively, the high affinity CH848 trimer exhibited a strongly favorable binding entropy (-TΔS < 0) that is offset by a positive binding enthalpy (ΔH > 0) and a large, negative ΔC_p_. These data suggest that extensive conformational changes occur during association resulting in the slow association and dissociation rates observed for the highest affinity trimer. (**B**) A similar thermodynamic analysis of CH31 binding to the GT6, GT8, and A244 monomeric antigens also suggests varying binding mechanisms. The moderately fast on-rate, weak affinity GT6 (k_a_ = 4.0x10^4^ M^-1^s^-1^, k_d_ = 2.1x10^-1^ s^-1^, K_d_ = 5247 nM) demonstrates a strongly favorable binding enthalpy (ΔH << 0) that is offset by an unfavorable binding entropy (-TΔS > 0), whereas the fast on-rate, moderate affinity GT8 (k_a_ = 2.3x10^5^ M^-1^s^-1^, k_d_ = 7.0x10^-2^ s^-1^, K_d_ = 307.8 nM) demonstrates a favorable binding enthalpy (ΔH < 0) and a favorable binding entropy (-TΔS < 0). Both antigens bind with small ΔC_p_ values indicating that minimal conformational changes occur during binding. In contrast, despite a similar thermodynamic profile to GT8, the slow on-rate, high affinity A244 antigen (k_a_ = 2.4x10^3^ M^-1^s^-1^, k_d_ = 7.3x10^-5^ s^-1^, K_d_ = 30.5 nM) exhibits a strongly negative ΔC_p_ indicating extensive conformational change is required to facilitate the bound complex. The observed relationship between thermodynamic conformational hurdles and antigen association kinetics provides further support to a kinetic model for B cell activation. Values and error bars represent the average and standard deviation of three measurements.

**Figure S6:**
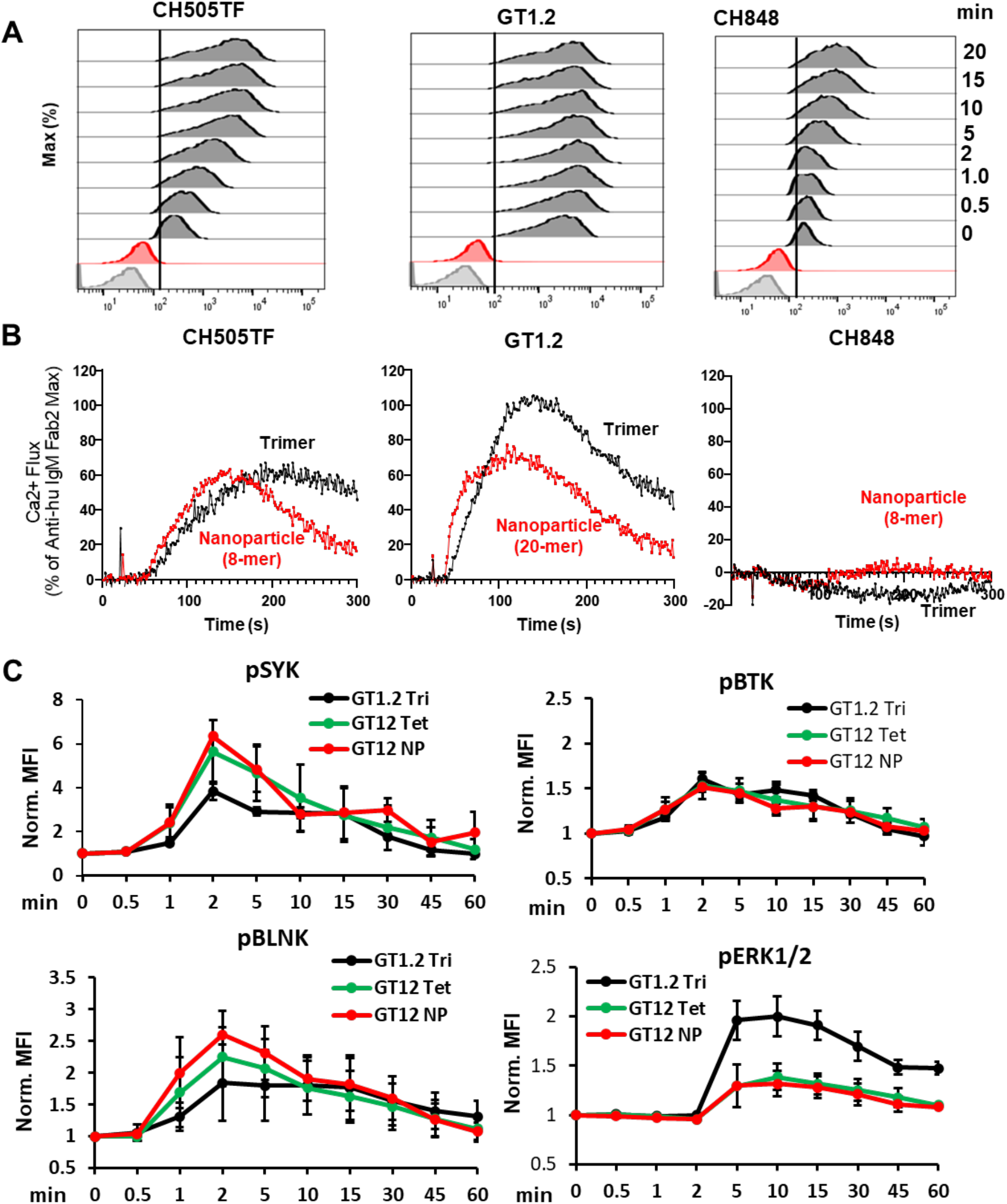
Cell surface binding and signaling induced by SOSIP trimers and multimers of trimers. (**A**) Representative flow plots showing binding of CH505TF, GT1.2 and CH848 trimers to CH31 IgM BCRs over a period of 20 mins. Cells were stimulated with the biotinylated trimeric antigens at 30 nM concentration at 37°C and aliquots of cells were collected at the indicated timepoints which further fixed and stained by anti-biotin antibody to analyzed in flow cytometer. (**B**) Comparison of the CH31 IgM Ramos cell activation induced by three trimeric antigens versus their nanoparticle forms. Results include (from left to right) CH505TF trimer (black) compared with the CH505TF ferritin nanoparticle (8-mer) (red); the calcium flux induced by the GT1.2 Trimer (black) compared with the GT1.2 nanoparticle (20-mer) (red); and the signaling mediated by the CH848 trimer (black) compared with its ferritin nanoparticle (8-mer) (red). Trimer and nanoparticle antigens for B cell activation were used at the same per unit trimer concentrations. Results are presented as a % of the maximum Anti-human IgM F(ab)2 response and are representative of at least two measurements. (**C**) B cell signaling kinetics by multimers of GT1.2 trimer antigen stimulation. The kinetics of proximal Syk, BLNK, Btk and ERK1/2 phosphorylation in CH31 IgM Ramos cells were measured over an hour of stimulation by GT1.2 trimer (black), tetramer of GT1.2 trimer (green) and 20-mer of GT1.2 trimer (red) at 30 nM concentration. Around 2.5x 10^5^ cells were collected at each indicated period of times, fixed (30 mins), permeabilized and then stained for phosphorylated Syk, BLNK, Btk and ERK1/2 and analyzed by BD LSRII flow cytometer. Plots shown here is normalized MFI data from three independent experiments. At each time point, the phosphorylation was normalized to the unstimulated cells MFI value (normalized value =1 for the unstimulated, unphosphorylated subset). Error bar indicate standard deviation of means. **Methods**. Flow analysis was performed as described in the methods section “Antigen binding analysis in CH31 IgM Ramos cells”. Calcium flux experiments were performed as described in the methods section “Calcium Flux Measurements with Ramos cells”; however, due to assay limitations, the concentrations of the trimers and nanoparticles were lower than the previously described 1uM. The final concentrations of the trimers and nanoparticles of CH505TF and CH848 were 0.5uM while the GT1.2 trimer and nanoparticle were compared at 0.15uM. B cell signaling by multimers of GT1.2 trimer was described in the method section “Phospho-flow kinase analysis in CH31 IgM Ramos”.

**Figure S7.**
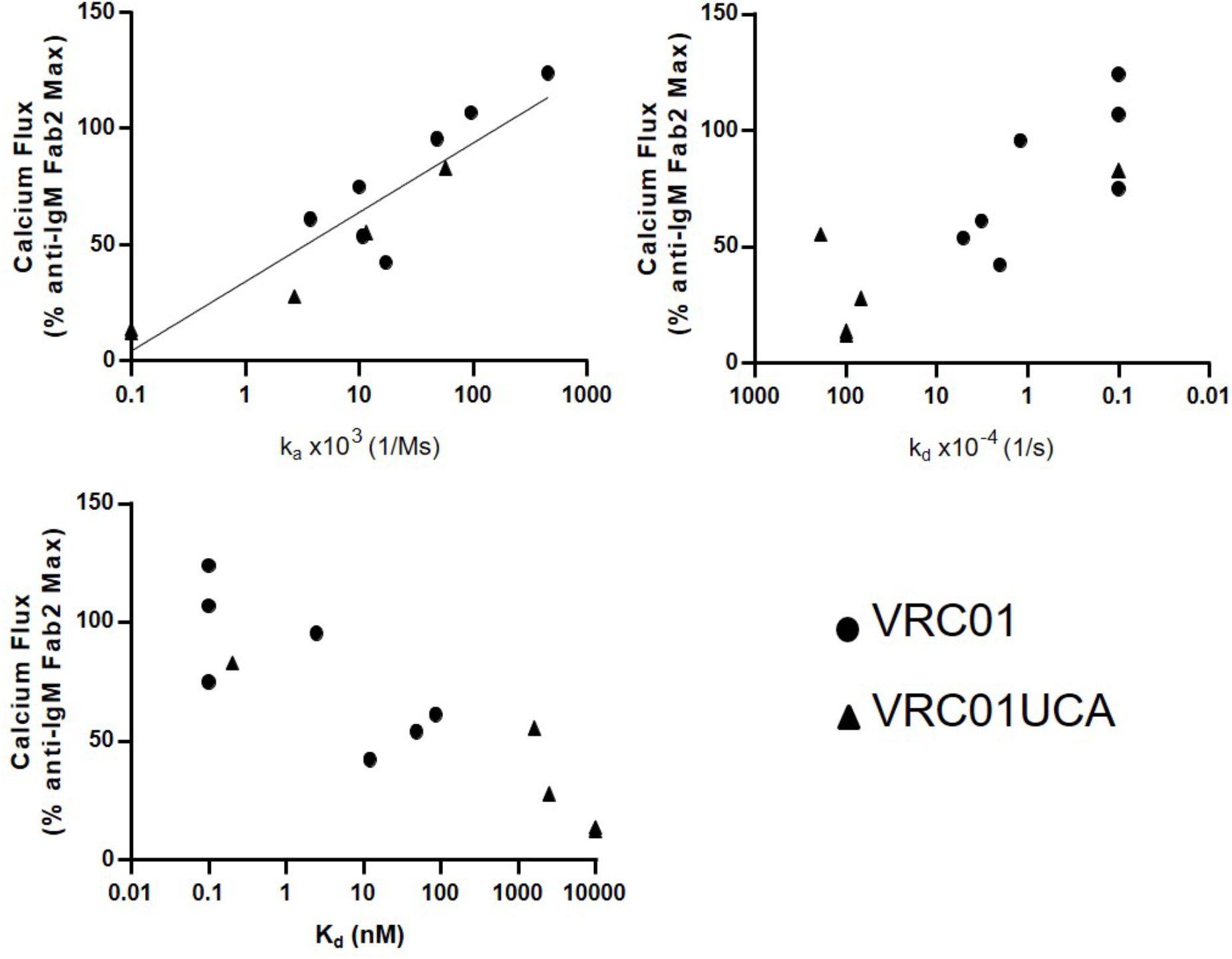
Relationship between calcium flux response and antigen binding kinetic rates for VRC01 and VRC01UCA Ramos cell lines. VRC01 and VRC01UCA IgM Ramos cell calcium flux mediated by multimerized antigens (y-axis) plotted versus SPR measured monomeric (or trimeric) antigen on-rate (k_a_, 1/Ms) (top left), off-rate (k_d_, 1/s) (top right) or affinity (K_d_, nM) (bottom left) against VRC01 or VRC01UCA bnAb (x-axis). Calcium flux responses are presented as a percentage of the maximum Anti-human IgM F(ab’)2 mediated responses and both calcium flux and kinetic rate parameters are representative one measurement for VRC01/VRC01UCA results. Correlation evaluation was assessed by Kendall’s Tau analysis (k_a_ and VRC01 UCA: Kendall’s Tau 0.9487; p-value = 0.0230; _d_ and VRC01 UCA: Kendall’s Tau - 0.9487; p-value = 0.0230). The association between K_d_ and VRCO1UCA IgM Ca-flux did not reach significance (K_d_ and VRC01: Kendall’s Tau -0.6172; p-value = 0.0599). **Methods:** Calcium flux experiments were performed as described in the methods section “Calcium Flux Measurements with Ramos cells”. SPR affinities and kinetic rates were measured as previously described (Bonsignori, et al., 2018). To assess potential correlations, Kendall’s Tau was used due to the small sample size. The alpha level was set at 0.05 and there were no adjustments made for multiple correlation evaluations. The analysis was conducted using SAS 9.4 software.

**Figure S8.**
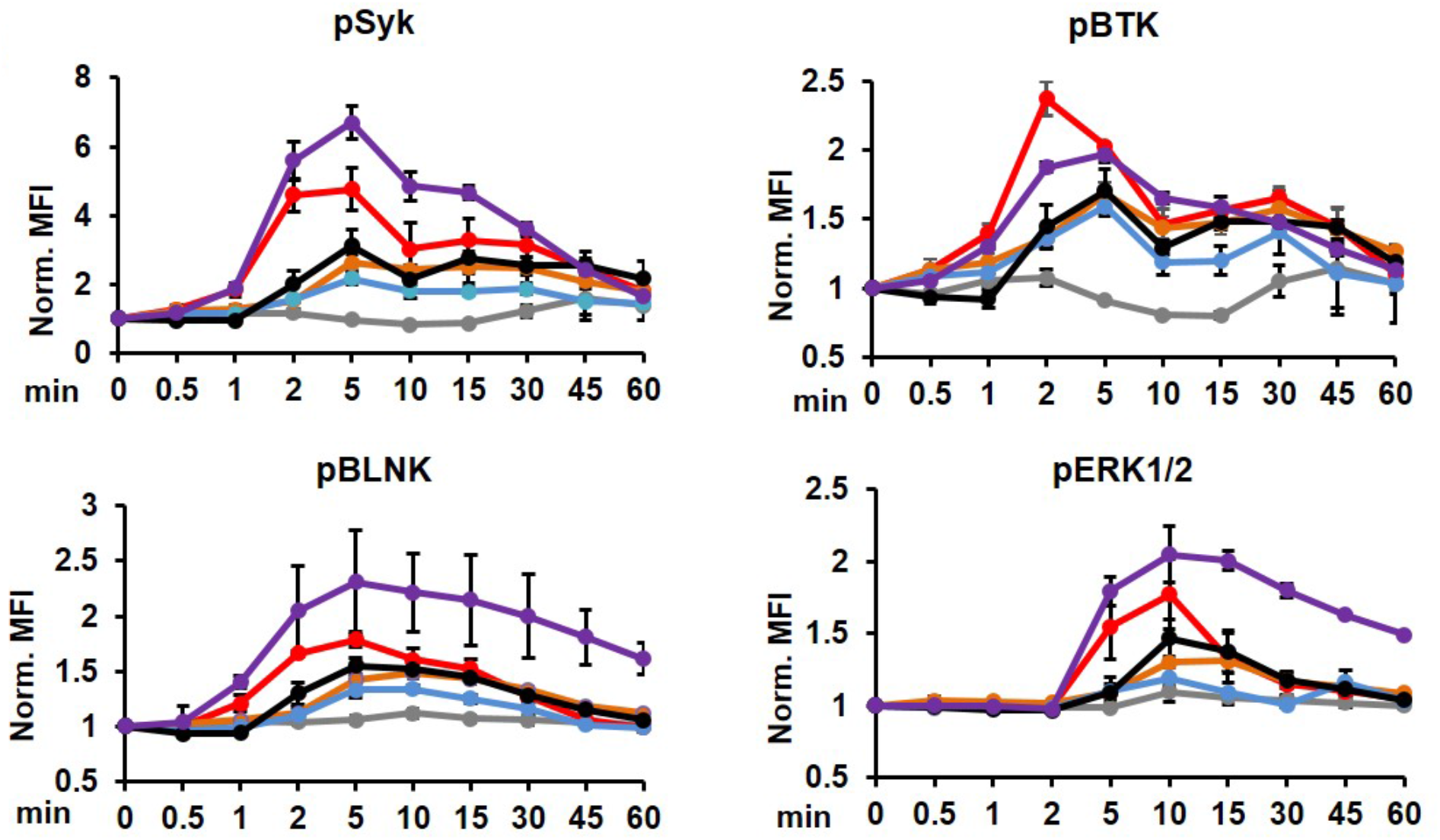
CH31 IgM B phospho signaling by tetrameric antigens. B cell signaling kinetics by tetramers of gp120 based antigen stimulation. The kinetics of proximal Syk, BLNK, Btk and ERK1/2 phosphorylation in CH31 IgM Ramos cells were measured over an hour of stimulation by GT6 (grey), GT8 (red), A244 (orange), CH505TF (blue) and 426c (black) at 250 nM concentration along with anti-human IgM (purple) at 30 nM concentration. Around 2.5x10^5^ cells were collected at each indicated period of times, fixed (30 mins), permeabilized and then stained for phosphorylated Syk, BLNK, Btk and ERK1/2. Plots shown here is normalized MFI data from three independent experiments. At each time point, the phosphorylation level was normalized to the unstimulated cells MFI value (normalized value =1 for the unstimulated, unphosphorylated subset). Error bars indicate standard deviation of means. **Methods:** Experimental conditions are as described in the methods section under “Antigen binding analysis in CH31 IgM Ramos cells” and “Phospho-flow kinase analysis in CH31 IgM Ramos”.

**Figure S9.**
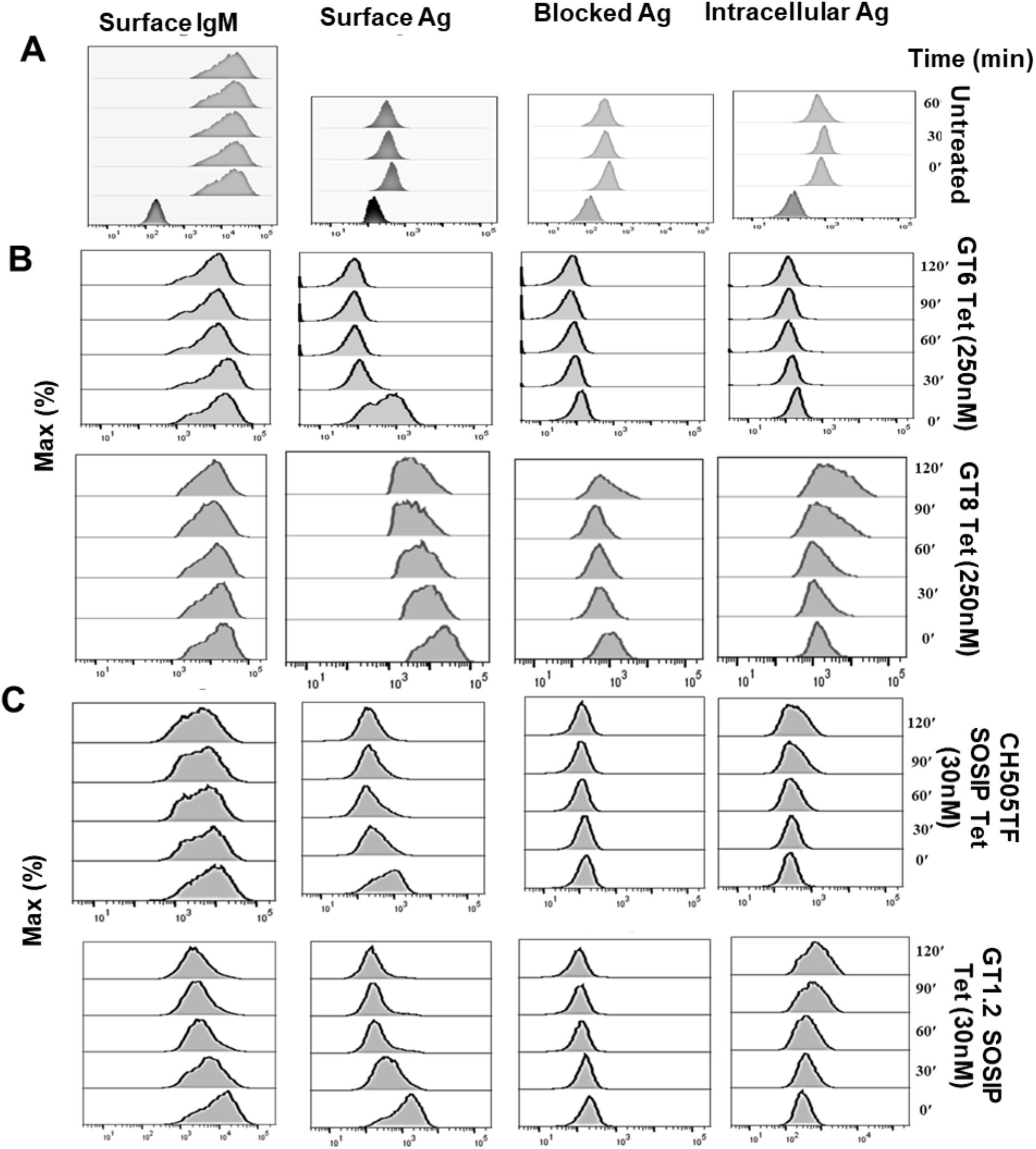
Flow cytometric detection of intracellular antigens in CH31 IgM Ramos cells. Representative flow plots showing level of surface IgM, surface antigens, blocked antigens and intracellular antigens detected over time in ramos cells incubated with SA-conjugated tetramers of gp120 based antigens **(B)** and tetramer of trimeric antigens **(C)** or left untreated **(A)**. Level of surface IgM and bound antigens were detected at each time points using flourophore conjugated anti-IgM and anti-SA antibody respectively. To block the SA-binding sites on surface antigens, cells were incubated with non-fluorophore anti-SA antibody and blocked level of SA binding sites were confirmed. For intracellular antigen detection, cells were fixed and permeabilized after surface blockage of SA binding sites and further incubated with fluorophore conjugated anti-SA antibody and analyzed by flow cytometry. **Methods:** Experiments were performed as described in the methods section under “Antigen internalization assay in CH31 IgM Ramos”.

**Figure S10.**
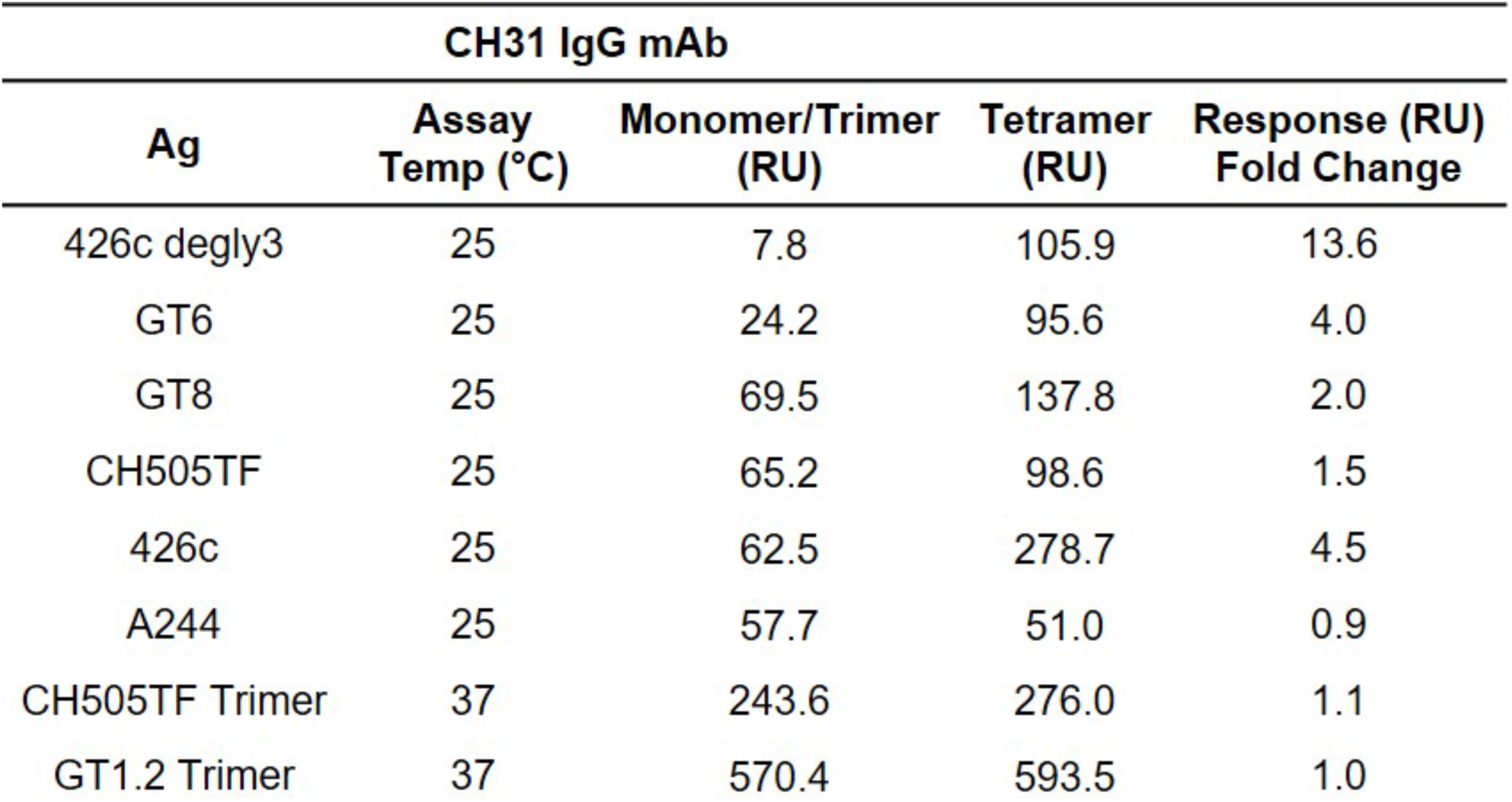
Response fold change of antigen monomers/trimers versus multimers. Summary table comparing the SPR binding responses (RU) to CH31 IgG bnAb of monomeric antigens (or trimeric) with the binding responses of their multimeric (4-mer) forms. The responses were measured at the end of the association phase of the sample injection and the fold change between the antigen types was calculated by dividing the response of the tetramer by the response of the monomer (or trimer). The 426c degly3 and 426c core proteins exhibited the largest increase in binding response against CH31 IgG after tetramerization. GT6, GT8, and CH505TF gp120 all showed some enhancement in binding (1.5x or greater) after tetramerization while A244, CH505TF trimer, and GT1.2 trimer showed little to no increase in binding. **Methods:** SPR experiments with monomeric (or trimeric) antigens versus low-order multimers of the same antigens were performed as described in the methods section under “Surface Plasmon Resonance of Tetrameric Antigens”.

## References

1. Abbott, R.K., Lee, J.H., Menis, S., Skog, P., Rossi, M., Ota, T., Kulp, D.W., Bhullar, D., Kalyuzhniy, O., Havenar-Daughton, C., et al. (2018). Precursor Frequency and Affinity Determine B Cell Competitive Fitness in Germinal Centers, Tested with Germline-Targeting HIV Vaccine Immunogens. Immunity 48, 133–146 e136.

2. Alam, S.M., Liao, H.X., Tomaras, G.D., Bonsignori, M., Tsao, C.Y., Hwang, K.K., Chen, H., Lloyd, K.E., Bowman, C., Sutherland, L., et al. (2013). Antigenicity and immunogenicity of RV144 vaccine AIDSVAX clade E envelope immunogen is enhanced by a gp120 N-terminal deletion. J Virol 87, 1554–1568.

3. Batista, F.D., and Neuberger, M.S. (1998). Affinity dependence of the B cell response to antigen: a threshold, a ceiling, and the importance of off-rate. Immunity 8, 751–759.

4. Bonsignori, M., Liao, H.X., Gao, F., Williams, W.B., Alam, S.M., Montefiori, D.C., and Haynes, B.F. (2017). Antibody-virus co-evolution in HIV infection: paths for HIV vaccine development. Immunol Rev 275, 145–160.

5. Bonsignori, M., Montefiori, D.C., Wu, X., Chen, X., Hwang, K.K., Tsao, C.Y., Kozink, D.M., Parks, R.J., Tomaras, G.D., Crump, J.A., et al. (2012). Two distinct broadly neutralizing antibody specificities of different clonal lineages in a single HIV-1-infected donor: implications for vaccine design. J Virol 86, 4688–4692.

6. Bonsignori, M., Scott, E., Wiehe, K., Easterhoff, D., Alam, S.M., Hwang, K.K., Cooper, M., Xia, S.M., Zhang, R., Montefiori, D.C., et al. (2018). Inference of the HIV-1 VRC01 Antibody Lineage Unmutated Common Ancestor Reveals Alternative Pathways to Overcome a Key Glycan Barrier. Immunity 49, 1162–1174 e1168.

7. Brouwer, P.J.M., and Sanders, R.W. (2019). Presentation of HIV-1 envelope glycoprotein trimers on diverse nanoparticle platforms. Curr Opin HIV AIDS 14, 302–308.

8. Chan, T.D., and Brink, R. (2012). Affinity-based selection and the germinal center response. Immunol Rev 247, 11–23.

9. Chen, Y., Zhang, J., Hwang, K.K., Bouton-Verville, H., Xia, S.M., Newman, A., Ouyang, Y.B., Haynes, B.F., and Verkoczy, L. (2013). Common tolerance mechanisms, but distinct cross-reactivities associated with gp41 and lipids, limit production of HIV-1 broad neutralizing antibodies 2F5 and 4E10. J Immunol 191, 1260–1275.

10. Dal Porto, J.M., Haberman, A.M., Kelsoe, G., and Shlomchik, M.J. (2002). Very low affinity B cells form germinal centers, become memory B cells, and participate in secondary immune responses when higher affinity competition is reduced. J Exp Med 195, 1215–1221.

11. Dennison, S.M., Sutherland, L.L., Jaeger, F.H., Anasti, K.M., Parks, R., Stewart, S., Bowman, C., Xia, S.M., Zhang, R., Shen, X., et al. (2011). Induction of antibodies in rhesus macaques that recognize a fusion-intermediate conformation of HIV-1 gp41. PLoS One 6, e27824.

12. Dintzis, R.Z., Vogelstein, B., and Dintzis, H.M. (1982). Specific cellular stimulation in the primary immune response: experimental test of a quantized model. Proc Natl Acad Sci U S A 79, 884–888.

13. Dosenovic, P., Kara, E.E., Pettersson, A.K., McGuire, A.T., Gray, M., Hartweger, H., Thientosapol, E.S., Stamatatos, L., and Nussenzweig, M.C. (2018). Anti-HIV-1 B cell responses are dependent on B cell precursor frequency and antigen-binding affinity. Proc Natl Acad Sci U S A 115, 4743–4748.

14. Fairhead, M., Krndija, D., Lowe, E.D., and Howarth, M. (2014). Plug-and-play pairing via defined divalent streptavidins. J Mol Biol 426, 199–214.

15. Foote, J., and Eisen, H.N. (1995). Kinetic and affinity limits on antibodies produced during immune responses. Proc Natl Acad Sci U S A 92, 1254–1256.

16. Foote, J., and Eisen, H.N. (2000). Breaking the affinity ceiling for antibodies and T cell receptors. Proc Natl Acad Sci U S A 97, 10679–10681.

17. Haynes, B.F., and Mascola, J.R. (2017). The quest for an antibody-based HIV vaccine. Immunol Rev 275, 5–10.

18. Henderson, R., Watts, B.E., Ergin, H.N., Anasti, K., Parks, R., Xia, S.M., Trama, A., Liao, H.X., Saunders, K.O., Bonsignori, M., et al. (2019). Selection of immunoglobulin elbow region mutations impacts interdomain conformational flexibility in HIV-1 broadly neutralizing antibodies. Nat Commun 10, 654.

19. Iype, J., Datta, M., Khadour, A., Ubelhart, R., Nicolo, A., Rollenske, T., Duhren-von Minden, M., Wardemann, H., Maity, P.C., and Jumaa, H. (2019). Differences in Self-Recognition between Secreted Antibody and Membrane-Bound B Cell Antigen Receptor. J Immunol 202, 1417–1427.

20. Jardine, J., Julien, J.P., Menis, S., Ota, T., Kalyuzhniy, O., McGuire, A., Sok, D., Huang, P.S., MacPherson, S., Jones, M., et al. (2013). Rational HIV immunogen design to target specific germline B cell receptors. Science 340, 711–716.

21. Jiang, A., Craxton, A., Kurosaki, T., and Clark, E.A. (1998). Different protein tyrosine kinases are required for B cell antigen receptor-mediated activation of extracellular signal-regulated kinase, c-Jun NH2-terminal kinase 1, and p38 mitogen-activated protein kinase. J Exp Med 188, 1297–1306.

22. Jumaa, H., Hendriks, R.W., and Reth, M. (2005). B cell signaling and tumorigenesis. Annu Rev Immunol 23, 415–445.

23. Kepler, T.B., Liao, H.X., Alam, S.M., Bhaskarabhatla, R., Zhang, R., Yandava, C., Stewart, S., Anasti, K., Kelsoe, G., Parks, R., et al. (2014). Immunoglobulin gene insertions and deletions in the affinity maturation of HIV-1 broadly reactive neutralizing antibodies. Cell Host Microbe 16, 304–313.

24. Kouskoff, V., Famiglietti, S., Lacaud, G., Lang, P., Rider, J.E., Kay, B.K., Cambier, J.C., and Nemazee, D. (1998). Antigens varying in affinity for the B cell receptor induce differential B lymphocyte responses. J Exp Med 188, 1453–1464.

25. Krogsgaard, M., Prado, N., Adams, E.J., He, X.L., Chow, D.C., Wilson, D.B., Garcia, K.C., and Davis, M.M. (2003). Evidence that structural rearrangements and/or flexibility during TCR binding can contribute to T cell activation. Mol Cell 12, 1367–1378.

26. Kuraoka, M., Schmidt, A.G., Nojima, T., Feng, F., Watanabe, A., Kitamura, D., Harrison, S.C., Kepler, T.B., and Kelsoe, G. (2016). Complex Antigens Drive Permissive Clonal Selection in Germinal Centers. Immunity 44, 542–552.

27. Kurosaki, T. (2002). Regulation of B-cell signal transduction by adaptor proteins. Nat Rev Immunol 2, 354–363.

28. Lane, P.J., Ledbetter, J.A., McConnell, F.M., Draves, K., Deans, J., Schieven, G.L., and Clark, E.A. (1991). The role of tyrosine phosphorylation in signal transduction through surface Ig in human B cells. Inhibition of tyrosine phosphorylation prevents intracellular calcium release. J Immunol 146, 715–722.

29. Liao, H.X., Lynch, R., Zhou, T., Gao, F., Alam, S.M., Boyd, S.D., Fire, A.Z., Roskin, K.M., Schramm, C.A., Zhang, Z., et al. (2013). Co-evolution of a broadly neutralizing HIV-1 antibody and founder virus. Nature 496, 469–476.

30. Maity, P.C., Blount, A., Jumaa, H., Ronneberger, O., Lillemeier, B.F., and Reth, M. (2015). B cell antigen receptors of the IgM and IgD classes are clustered in different protein islands that are altered during B cell activation. Sci Signal 8, ra93.

31. McGuire, A.T., Gray, M.D., Dosenovic, P., Gitlin, A.D., Freund, N.T., Petersen, J., Correnti, C., Johnsen, W., Kegel, R., Stuart, A.B., et al. (2016). Specifically modified Env immunogens activate B-cell precursors of broadly neutralizing HIV-1 antibodies in transgenic mice. Nat Commun 7, 10618.

32. McGuire, A.T., Hoot, S., Dreyer, A.M., Lippy, A., Stuart, A., Cohen, K.W., Jardine, J., Menis, S., Scheid, J.F., West, A.P., et al. (2013). Engineering HIV envelope protein to activate germline B cell receptors of broadly neutralizing anti-CD4 binding site antibodies. J Exp Med 210, 655–663.

33. Mizuno, T., and Rothstein, T.L. (2005). B cell receptor (BCR) cross-talk: CD40 engagement enhances BCR-induced ERK activation. J Immunol 174, 3369–3376.

34. Packard, T.A., and Cambier, J.C. (2013). B lymphocyte antigen receptor signaling: initiation, amplification, and regulation. F1000Prime Rep 5, 40.

35. Paus, D., Phan, T.G., Chan, T.D., Gardam, S., Basten, A., and Brink, R. (2006). Antigen recognition strength regulates the choice between extrafollicular plasma cell and germinal center B cell differentiation. J Exp Med 203, 1081–1091.

36. Phan, T.G., Paus, D., Chan, T.D., Turner, M.L., Nutt, S.L., Basten, A., and Brink, R. (2006). High affinity germinal center B cells are actively selected into the plasma cell compartment. J Exp Med 203, 2419–2424.

37. Pierce, S.K., and Liu, W. (2010). The tipping points in the initiation of B cell signalling: how small changes make big differences. Nat Rev Immunol 10, 767–777.

38. Pugach, P., Ozorowski, G., Cupo, A., Ringe, R., Yasmeen, A., de Val, N., Derking, R., Kim, H.J., Korzun, J., Golabek, M., et al. (2015). A native-like SOSIP.664 trimer based on an HIV-1 subtype B env gene. J Virol 89, 3380–3395.

39. Reth, M., and Wienands, J. (1997). Initiation and processing of signals from the B cell antigen receptor. Annu Rev Immunol 15, 453–479.

40. Sanders, R.W., van Gils, M.J., Derking, R., Sok, D., Ketas, T.J., Burger, J.A., Ozorowski, G., Cupo, A., Simonich, C., Goo, L., et al. (2015). HIV-1 VACCINES. HIV-1 neutralizing antibodies induced by native-like envelope trimers. Science 349, aac4223.

41. Saunders, K.O., Verkoczy, L.K., Jiang, C., Zhang, J., Parks, R., Chen, H., Housman, M., Bouton-Verville, H., Shen, X., Trama, A.M., et al. (2017). Vaccine Induction of Heterologous Tier 2 HIV-1 Neutralizing Antibodies in Animal Models. Cell Rep 21, 3681–3690.

42. Saunders, K.O., Wiehe, K., Tian, M., Acharya, P., Bradley, T., Alam, S.M., Go, E.P., Scearce, R., Sutherland, L., Henderson, R., et al. (2019). Targeted selection of HIV-specific antibody mutations by engineering B cell maturation. Science 366.

43. Schamel, W.W., and Reth, M. (2000). Monomeric and oligomeric complexes of the B cell antigen receptor. Immunity 13, 5–14.

44. Sela-Culang, I., Alon, S., and Ofran, Y. (2012). A systematic comparison of free and bound antibodies reveals binding-related conformational changes. J Immunol 189, 4890–4899.

45. Shih, T.A., Meffre, E., Roederer, M., and Nussenzweig, M.C. (2002). Role of BCR affinity in T cell dependent antibody responses in vivo. Nat Immunol 3, 570–575.

46. Sulzer, B., and Perelson, A.S. (1997). Immunons revisited: binding of multivalent antigens to B cells. Mol Immunol 34, 63–74.

47. Takata, M., Sabe, H., Hata, A., Inazu, T., Homma, Y., Nukada, T., Yamamura, H., and Kurosaki, T. (1994). Tyrosine kinases Lyn and Syk regulate B cell receptor-coupled Ca2+ mobilization through distinct pathways. EMBO J 13, 1341–1349.

48. Tolar, P., Sohn, H.W., Liu, W., and Pierce, S.K. (2009). The molecular assembly and organization of signaling active B-cell receptor oligomers. Immunol Rev 232, 34–41.

49. Tolar, P., and Spillane, K.M. (2014). Force generation in B-cell synapses: mechanisms coupling B-cell receptor binding to antigen internalization and affinity discrimination. Adv Immunol 123, 69–100.

50. Veneziano, R., Moyer, T.J., Stone, M.B., Wamhoff, E.C., Read, B.J., Mukherjee, S., Shepherd, T.R., Das, J., Schief, W.R., Irvine, D.J., and Bathe, M. (2020). Role of nanoscale antigen organization on B-cell activation probed using DNA origami. Nat Nanotechnol 15, 716–723.

51. Verkoczy, L., Chen, Y., Bouton-Verville, H., Zhang, J., Diaz, M., Hutchinson, J., Ouyang, Y.B., Alam, S.M., Holl, T.M., Hwang, K.K., et al. (2011). Rescue of HIV-1 broad neutralizing antibody-expressing B cells in 2F5 VH x VL knockin mice reveals multiple tolerance controls. J Immunol 187, 3785–3797.

52. Verkoczy, L., Diaz, M., Holl, T.M., Ouyang, Y.B., Bouton-Verville, H., Alam, S.M., Liao, H.X., Kelsoe, G., and Haynes, B.F. (2010). Autoreactivity in an HIV-1 broadly reactive neutralizing antibody variable region heavy chain induces immunologic tolerance. Proc Natl Acad Sci U S A 107, 181–186.

53. Viant, C., Weymar, G.H.J., Escolano, A., Chen, S., Hartweger, H., Cipolla, M., Gazumyan, A., and Nussenzweig, M.C. (2020). Antibody Affinity Shapes the Choice between Memory and Germinal Center B Cell Fates. Cell 183, 1298–1311 e1211.

54. Viant, C., Wirthmiller, T., ElTanbouly, M.A., Chen, S.T., Cipolla, M., Ramos, V., Oliveira, T.Y., Stamatatos, L., and Nussenzweig, M.C. (2021). Germinal center-dependent and -independent memory B cells produced throughout the immune response. J Exp Med 218.

55. Vogelstein, B., Dintzis, R.Z., and Dintzis, H.M. (1982). Specific cellular stimulation in the primary immune response: a quantized model. Proc Natl Acad Sci U S A 79, 395–399.

56. Williams, W.B., Zhang, J., Jiang, C., Nicely, N.I., Fera, D., Luo, K., Moody, M.A., Liao, H.X., Alam, S.M., Kepler, T.B., et al. (2017). Initiation of HIV neutralizing B cell lineages with sequential envelope immunizations. Nat Commun 8, 1732.

57. Yang, J., and Reth, M. (2010a). The dissociation activation model of B cell antigen receptor triggering. FEBS Lett 584, 4872–4877.

58. Yang, J., and Reth, M. (2010b). Oligomeric organization of the B-cell antigen receptor on resting cells. Nature 467, 465–469.

59. Zhang, R., Verkoczy, L., Wiehe, K., Munir Alam, S., Nicely, N.I., Santra, S., Bradley, T., Pemble, C.W.t., Zhang, J., Gao, F., et al. (2016). Initiation of immune tolerance-controlled HIV gp41 neutralizing B cell lineages. Sci Transl Med 8, 336ra362.

