## supplemental for "B cells discriminate HIV-1 Envelope protein affinities by sensing antigen binding association rates"

### Supplemental Information

A

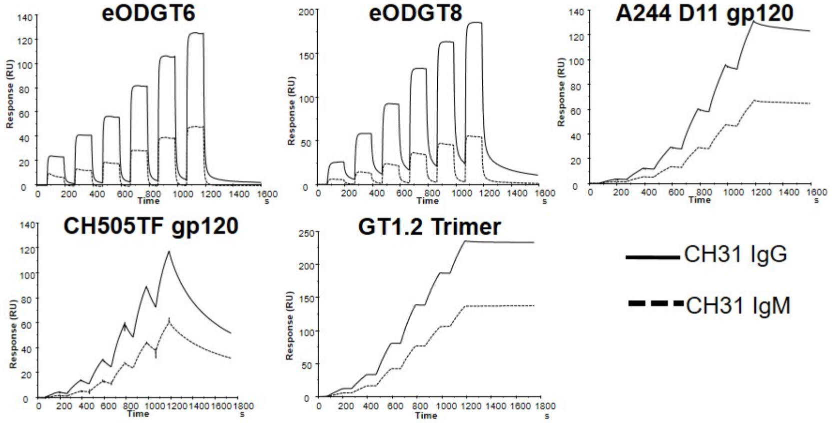


B

| Ag | Ab | $k_a$ (1/Ms) | $k_d$ (1/s) | $K_D$ (nM) |
|---|---|---|---|---|
| GT6 | CH31 IgG | 2.1E+4 | 1.2E-1 | 5,500 |
| | CH31 IgM | 2.2E+4 | 2.5E-1 | 11,000 |
| GT8 | CH31 IgG | 1.3E+5 | 1.2E-1 | 932 |
| | CH31 IgM | 2.0E+5 | 2.4E-1 | 1,222 |
| A244 | CH31 IgG | 1.4E+3 | 1.1E-4 | 78.7 |
| | CH31 IgM | 1.0E+3 | 7.0E-5 | 67.7 |
| CH505TF | CH31 IgG | 4.1E+3 | 7.1E-3 | 1,713 |
| | CH31 IgM | 1.5E+3 | 7.1E-3 | 4,743 |
| GT1.2 Trimer | CH31 IgG | 6.8E+4 | <1.0E-5 | <1.0 |
| | CH31 IgM | 5.5E+4 | <1.0E-5 | <1.0 |

**Figure S1. Affinity and kinetic rates of Env proteins binding to CH31 IgM mAb compared to CH31 IgG.** (**A**) SPR single cycle kinetic binding profiles of monomeric (or trimeric) antigens against CH31 IgG and CH31 IgM. Plots are representative of two measurements. (**B**) Summary of the overall binding affinities ($K_d$, nM) as well as the individual kinetic rate measurements ($k_a$ and $k_d$) of each antigen with either CH31 IgG or CH31 IgM. Values are representative of two data sets. A comparison of the binding results of the monomeric antigens (GT6, GT8, A244, and CH505TF) as well as the trimeric antigen, GT1.2, suggested that the affinities and kinetic rate parameters were similar for both isoforms of CH31.

**Methods:** SPR derived affinities and kinetic rates of the antigens against CH31 IgG and CH31 IgM were obtained using the Biacore S200 (Cytiva) instrument in HBS-EP+ 1X running buffer. CH31 IgG and CH31 IgM were directly immobilized onto two flow cells of a CM5 chip to a level of approximately 2000RU while a negative control antibody, Ab82, was immobilized onto flow cell 1 to a similar response level for reference subtraction. Six sequential injections of each antigen diluted from 5nM to 10,000nM were injected over the immobilized IgG and IgM mAbs at a flow rate of 50uL/min for 120s per injection. The single cycle injections were followed by a 600s dissociation and regeneration with a 20s pulse of Glycine pH2.0. The Biacore S200 Evaluation Software (Cytiva) was used to analyze results. Binding to the negative control antibody, Ab82, as well as buffer binding were used for double reference subtraction and accounted for both non-specific binding and signal drift. Curve fitting analyses were performed using the 1:1 Langmuir model with a local Rmax for all antigens except CH505TF. The CH505TF gp120 titration curve was fit using the heterogeneous ligand model with faster kinetic rate parameters reported. The reported kinetic rates and affinities are representative of 2 data sets.

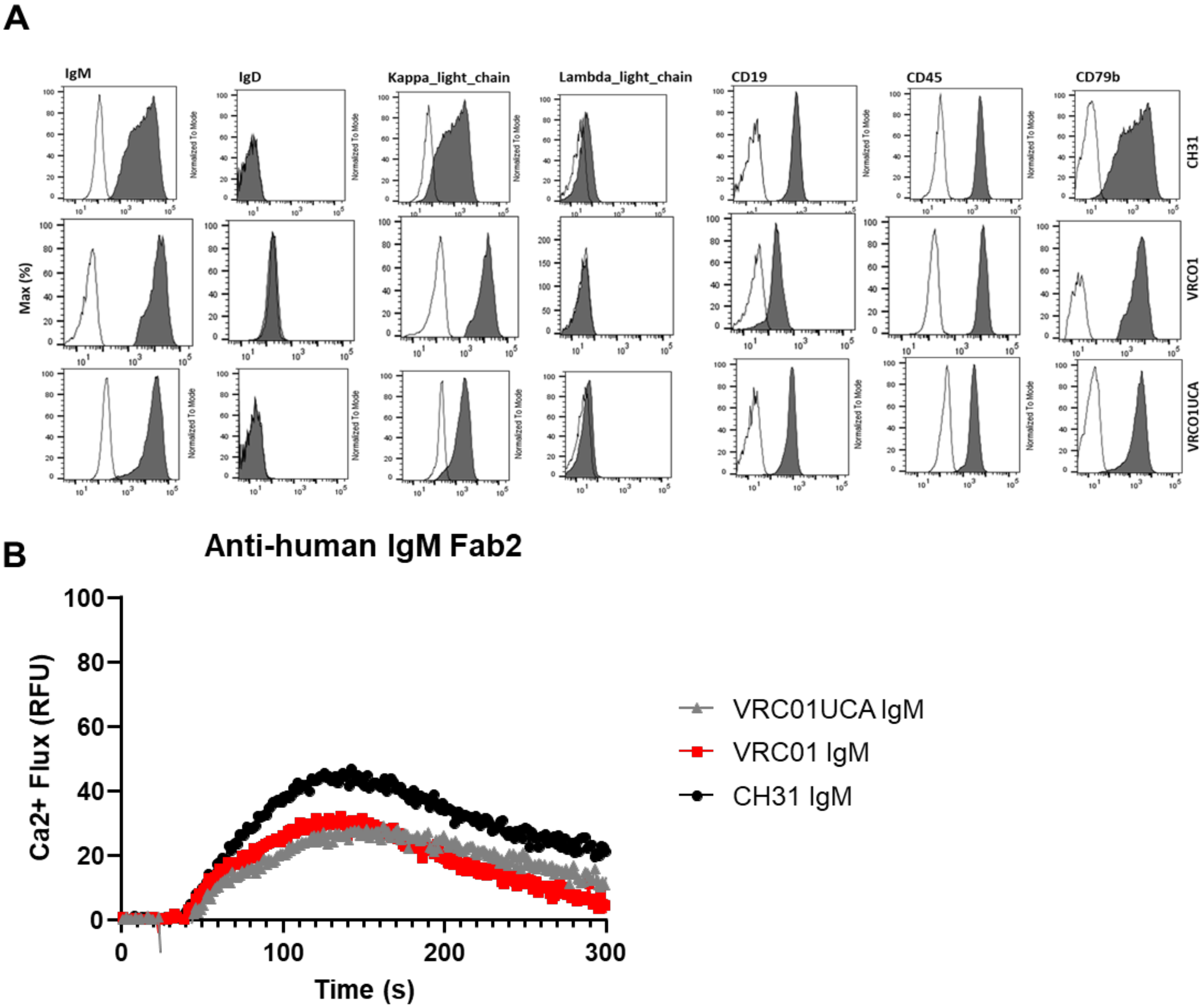


**Figure S2. Ramos cell line phenotypic data and activation with control stimulant**. (**A**) Representative flow cytometry plots showing expression of surface markers in bnAbs IgM (CH31, VRCO1 and VRCO1UCA) expressing ramos cells (dark) with unstained control (clear). Around 1.0x10$^{6}$ cells were fixed and stained for the surface expression of IgM, IgD, k-light chain, ʎ-light chain, CD19, CD45, CD79b using fluor conjugated antibody and analyzed by BD LSRII flow cytometer. (**B**) Calcium flux of the following 3 Ramos cells lines: VRC01UCA IgM (gray), VRC01 IgM (red), and CH31 IgM (black) by the positive control stimulant Anti-human IgM F(ab)'2 at 50ug/mL. The fluorescence of a blank well containing only

RPMI/dye was used for background subtraction.  The calcium flux profiles are representative of at least two measurements and demonstrate the functionality of the three cell lines.

**Methods:** Calcium flux experiments were performed as described in the methods section under “Calcium Flux Measurements with Ramos cells”.

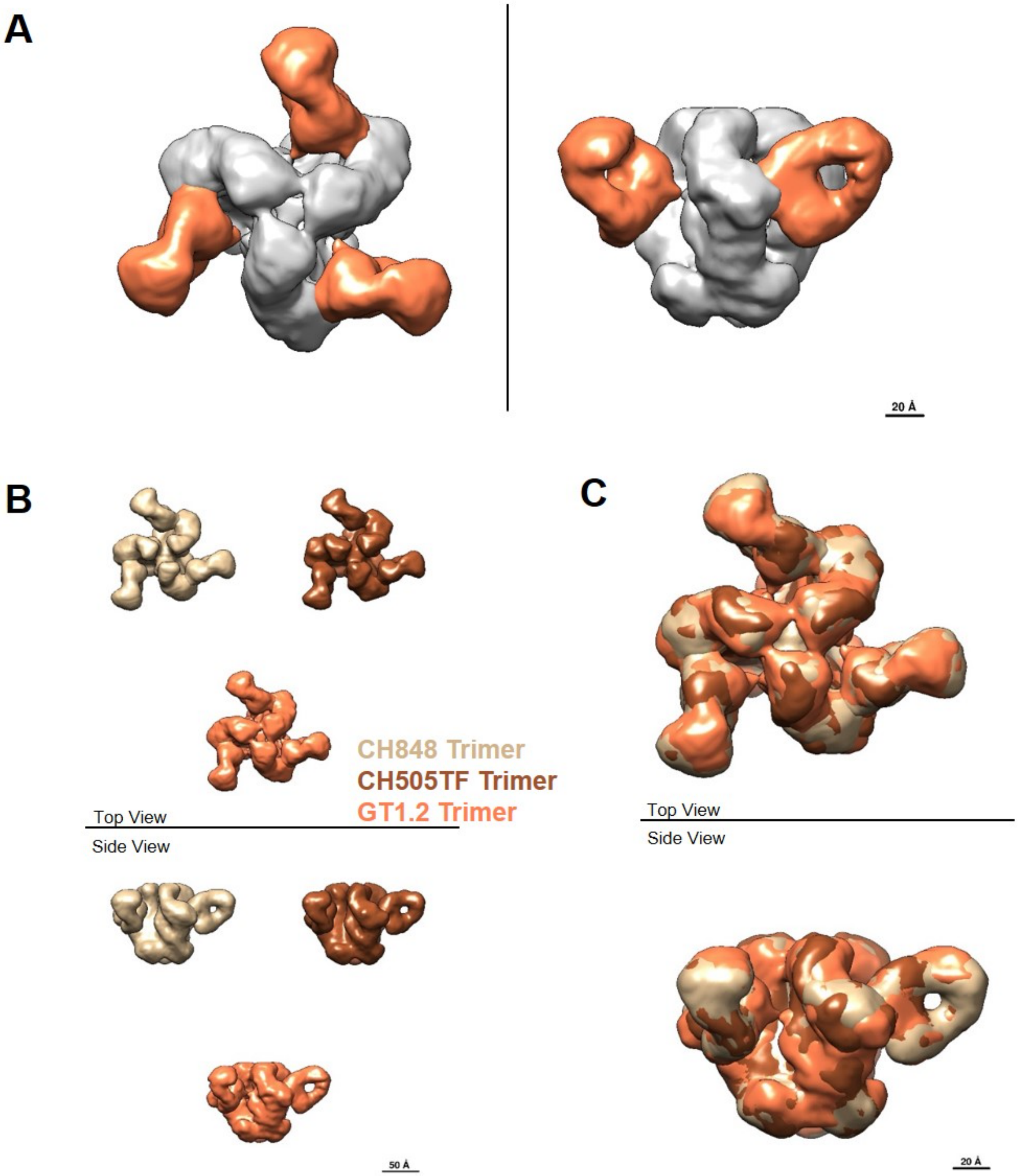


**Figure S3. Negative stain-EM structures of CH31 Fab complexed to three different SOSIP trimers.** (**A**) Structure of CH31 Fab bound to GT1.2 trimer, shown in top view and side view, with the density corresponding to the Env trimer colored gray and the density corresponding to the three Fabs colored orange. (**B**) Side by side comparison of the structures of CH31 Fab bound to CH848, CH505TF and GT1.2 trimers, colored tan, brown and orange, respectively. (**C**) Superposition of the three maps shown in **B** shows that at this resolution (~20 Å) there is no significant difference in the observed epitope or binding angle of CH31 to the three different SOSIP trimers.

**Methods:**

**Fab-SOSIP Complex Preparation and Purification**

Complex production was achieved by adding 15x Molar excess monoclonal Fab to 20ug trimer. This was then incubated overnight at room temperature. Fab-trimer complex was fixed by addition of glutaraldehyde to a final concentration of 5mM and then incubated for 10min at room temperature. Excess glutaraldehyde was quenched by adding sufficient 1M Tris and incubated for 10min at room temperature. Complex was purified through SEC by loading on a Superose 6 increase 10/300 column using a 100 µL loop, and run at 0.5 mL/min using an Äkta Pure system (GE Healthcare). Fractions were collected and complex peak pooled and concentrated by 10kDA cutoff centrifugal filtration (Amicon/EMD Millipore). Fab-SOSIP complex formation and quality was further accessed by SDS PAGE gel electrophoresis, where ~1 µg complex/lane was loaded on a 4%–15% TGX Stain free gel (BioRad) in both reducing and non-reducing conditions, and run at 200 V in Tris/Glycine/SDS buffer. Bands were visualized using Gel Doc EZ imager (BioRad), and the size of the fragments assessed by a protein standard ladder (BioRad).

**Negative stain electron microscopy**

Purified sample of each Fab-trimer complex was diluted to 100 µg/ml with HEPES-buffered saline (20 mM HEPES, 150 mM NaCl, pH 7.4) augmented with 5% glycerol and applied to a glow-discharged carbon-coated EM grid for 8-10 second. Sample was then blotted and stained with 2 g/dL uranyl formate for 1 min, blotted and air-dried. Grids were examined on a Philips EM420 electron microscope operating at 120 kV and nominal magnification of 49,000x, and 100-130 images were collected on a 76 Mpix CCD camera at 2.4 Å/pixel. Images were analyzed by 2D class averages and 3D reconstructions calculated using standard protocols with Relion 3.0 (Zivanov et al. 2018. *eLife*. 7:e42166).

A

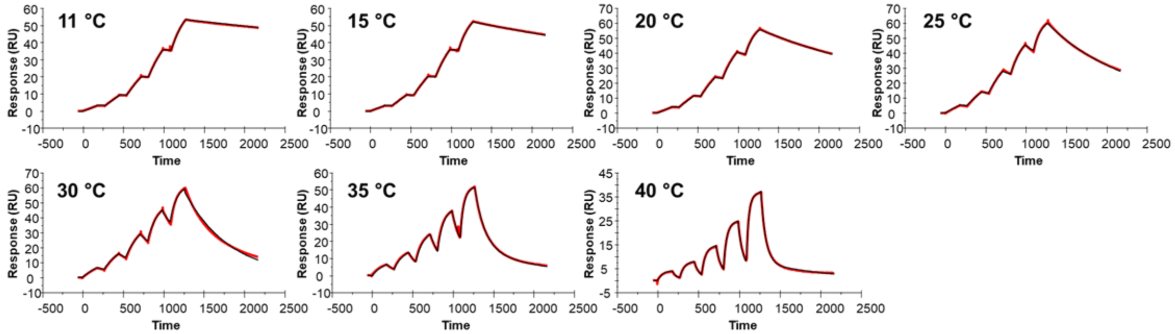


B

| Ab | Ag | Temp (°C) | $k_a$ (1/Ms) | $k_d$ (1/s) | $K_D$ (M) | $K_D$ (nM) |
|---|---|---|---|---|---|---|
| CH31 Fab | CH505TF | 11 | 2.92E+03 | 9.98E-05 | 3.42E-08 | 34.2 |
| CH31 Fab | CH505TF | 15 | 3.56E+03 | 1.80E-04 | 5.05E-08 | 50.5 |
| CH31 Fab | CH505TF | 20 | 4.83E+03 | 4.11E-04 | 8.52E-08 | 85.2 |
| CH31 Fab | CH505TF | 25 | 6.68E+03 | 9.98E-04 | 1.49E-07 | 149.3 |
| CH31 Fab | CH505TF | 30 | 1.05E+04 | 2.59E-03 | 2.47E-07 | 246.9 |
| CH31 Fab | CH505TF | 35 | 1.31E+04 | 8.12E-03 | 6.18E-07 | 617.8 |
| CH31 Fab | CH505TF | 40 | 1.42E+04 | 1.68E-02 | 1.18E-06 | 1181.6 |

C

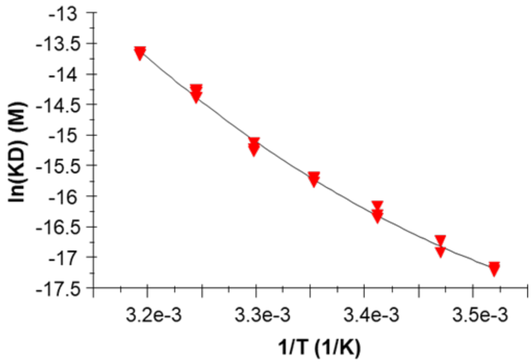


D

| *Parameter* | Value | SE |
|---|---|---|
| ΔH° [kJ/mol] | **-90.3** | 2.1 |
| ΔS° [J/(K*mol)] | **-170.0** | 10.0 |
| TΔS° [kJ/mol] | **-50.7** | 3.0 |
| -TΔS° [kJ/mol] | **50.7** | 3.0 |
| ΔG° [kJ/mol] | **-39.6** | 1.1 |
| ΔCp° [kJ/(K*mol)] | **-2.5** | 0.3 |

**Figure S4. Thermodynamic measurement of CH31 Fab binding to the CH505TF trimer by surface plasmon resonance (SPR).** (**A**) Single-cycle kinetic titration sensorgrams of CH31 Fab binding to the CH505TF SOSIP trimer recorded at varying temperatures. Plots are representative examples of 3 replicate measurements. (**B**) Table of association rate constants ($k_a$), dissociation rate constants ($k_d$), and affinity constants ($K_D$) recorded at varying temperatures. Values are reported as the average of 3 replicate measurements. (**C**) Plot of the natural logarithm of affinity ($K_D$) versus the reciprocal temperature and non-linear van't Hoff fit. (**D**) Table of thermodynamic parameters obtained from non-linear van't Hoff analysis. Values are reported as the average and standard deviation of 3 replicate measurements.

**Methods**: Thermodynamic analyses of CH31 binding to trimeric and monomeric antigens were performed by surface plasmon resonance (SPR) on a Biacore T200 platform (Cytiva Life Sciences) at varying temperatures in HBS-EP+ running buffer (10 mM HEPES, 150 mM NaCl, 3 mM EDTA, 0.05% Surfactant P20, pH 7.4). For trimeric antigens, biotinylated SOSIPs (50 µg/mL) were captured on streptavidin sensor surfaces at 5 µL/min to densities of 230-420 RU. Affinity measurements were performed by sequential titration of 5 concentrations of CH31 Fab [62.5-1000 nM (GT1.2), 100-1600 nM (CH505TF), 300-2400 nM (CH848)] over the SOSIP surfaces for 150-180 seconds per injection followed by a dissociation period of 900-2700 seconds at 30-50 µL/min. Surfaces were regenerated between Fab titrations with 10 mM glycine-HCl pH 2.0 for 40 seconds at 30-50 µL/min. Titrations were repeated for each trimer at a minimum of 7 different temperatures ranging from 11-45 °C. For monomeric antigens,

CH31 IgG mAb was captured on Protein A sensor surfaces at 5 µL/min to densities of 120-670 RU. Affinity measurements were performed by sequential titration of 5 concentrations of GT6 (625-10000 nM), GT8 (62.5-750 nM), or A244 (250-4000 nM) over a CH31 IgG surface for 180 seconds followed by a dissociation period of 720-1500 seconds at 50 µL/min. Surfaces were regenerated between antigen titrations with 10 mM glycine-HCl pH 1.5 for 45 seconds at 50 µL/min. Titrations were repeated for each antigen at 6 different temperatures ranging from 10-40 °C. Affinities were calculated using a steady state affinity model, a 1:1 kinetics model, or the fast components of a heterogeneous ligand kinetic model. Non-linear van't Hoff analyses were used to derive binding enthalpy (ΔH), entropy (ΔS), free energy (ΔG), and heat capacity ($\Delta C_p$).

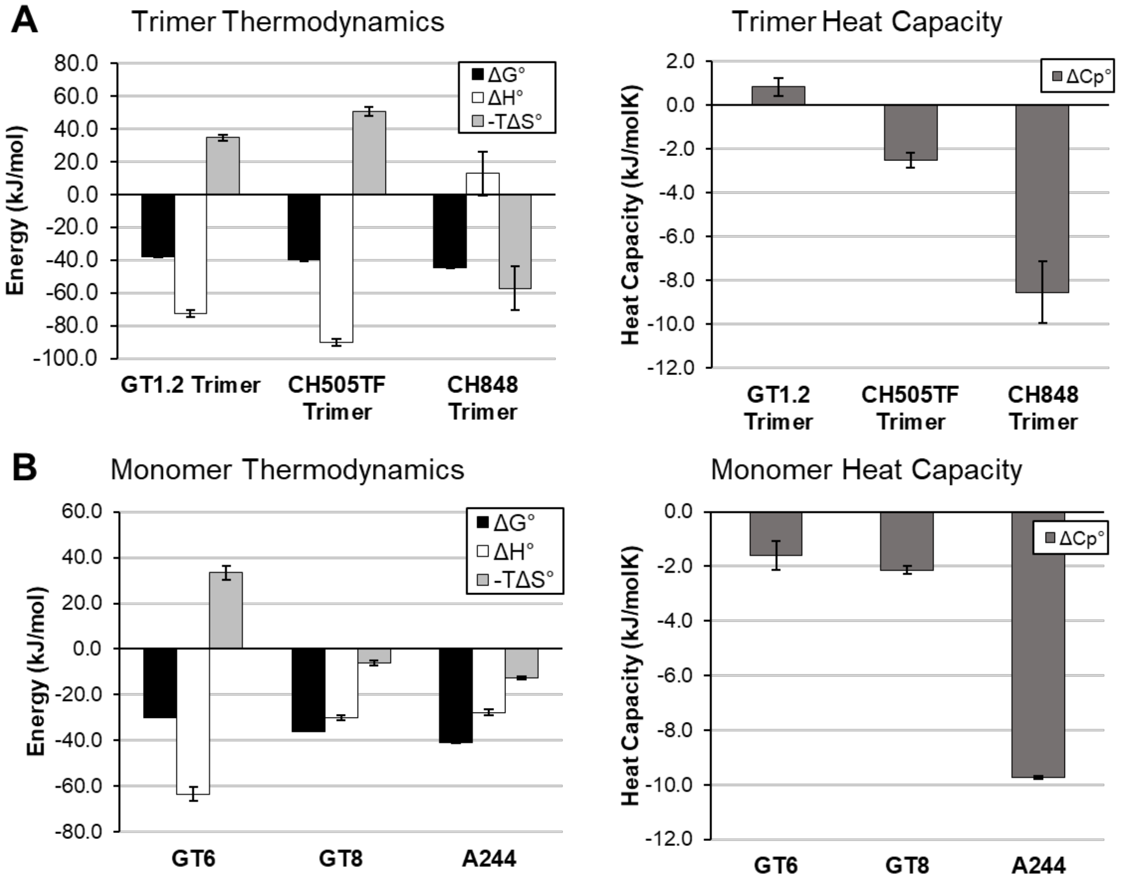


**Figure S5. Thermodynamics of CH31 binding to SOSIP trimers and monomeric antigens.** (**A**) Thermodynamic analysis of CH31 Fab binding to GT1.2, CH505TF, and CH848 trimers revealed markedly different binding profiles. The similar affinity GT1.2 and CH505TF trimers exhibited entropy-enthalpy compensation where the interaction was driven by strongly favorable binding enthalpies (ΔH < 0) and offset by unfavorable binding entropies (-TΔS > 0). The faster on-rate ($k_a = 2.2x10^5\ M^{-1}s^{-1}$) GT1.2 trimer displayed a relatively lower entropic hurdle (-TΔS) and a negligible $\Delta C_p$, which suggests that minimal conformational changes occur during the association process. Therefore, the GT1.2 trimer and CH31 Fab exist in unliganded states that require no conformational rearrangement to form a productive encounter complex. Alternatively, the high affinity CH848 trimer exhibited a strongly favorable binding entropy (-TΔS < 0) that is offset by a positive binding enthalpy (ΔH > 0) and a large, negative $\Delta C_p$. These data suggest that extensive conformational changes occur during association resulting in the slow association and dissociation rates observed for the highest affinity trimer. (**B**) A similar thermodynamic analysis of CH31 binding to the GT6, GT8, and A244 monomeric antigens also suggests varying binding mechanisms. The moderately fast on-rate, weak affinity GT6 ($k_a$ = 4.0x$10^4$ $M^{-1}s^{-1}$, $k_d$ = 2.1x$10^{-1}$ $s^{-1}$, $K_d$ = 5247 nM) demonstrates a strongly favorable binding enthalpy ($\Delta H \ll 0$) that is offset by an unfavorable binding entropy ($-T\Delta S > 0$), whereas the fast on-rate, moderate affinity GT8 ($k_a$ = 2.3x$10^5$ $M^{-1}s^{-1}$, $k_d$ = 7.0x$10^{-2}$ $s^{-1}$, $K_d$ = 307.8 nM) demonstrates a favorable binding enthalpy ($\Delta H < 0$) and a favorable binding entropy ($-T\Delta S < 0$). Both antigens bind with small $\Delta C_p$ values indicating that minimal conformational changes occur during binding. In contrast, despite a similar thermodynamic profile to GT8, the slow on-rate, high affinity A244 antigen ($k_a$ = 2.4x$10^3$ $M^{-1}s^{-1}$, $k_d$ = 7.3x$10^{-5}$ $s^{-1}$, $K_d$ = 30.5 nM) exhibits a strongly negative $\Delta C_p$ indicating extensive conformational change is required to facilitate the bound complex. The observed relationship between thermodynamic conformational hurdles and antigen association kinetics provides further support to a kinetic model for B cell activation. Values and error bars represent the average and standard deviation of three measurements.

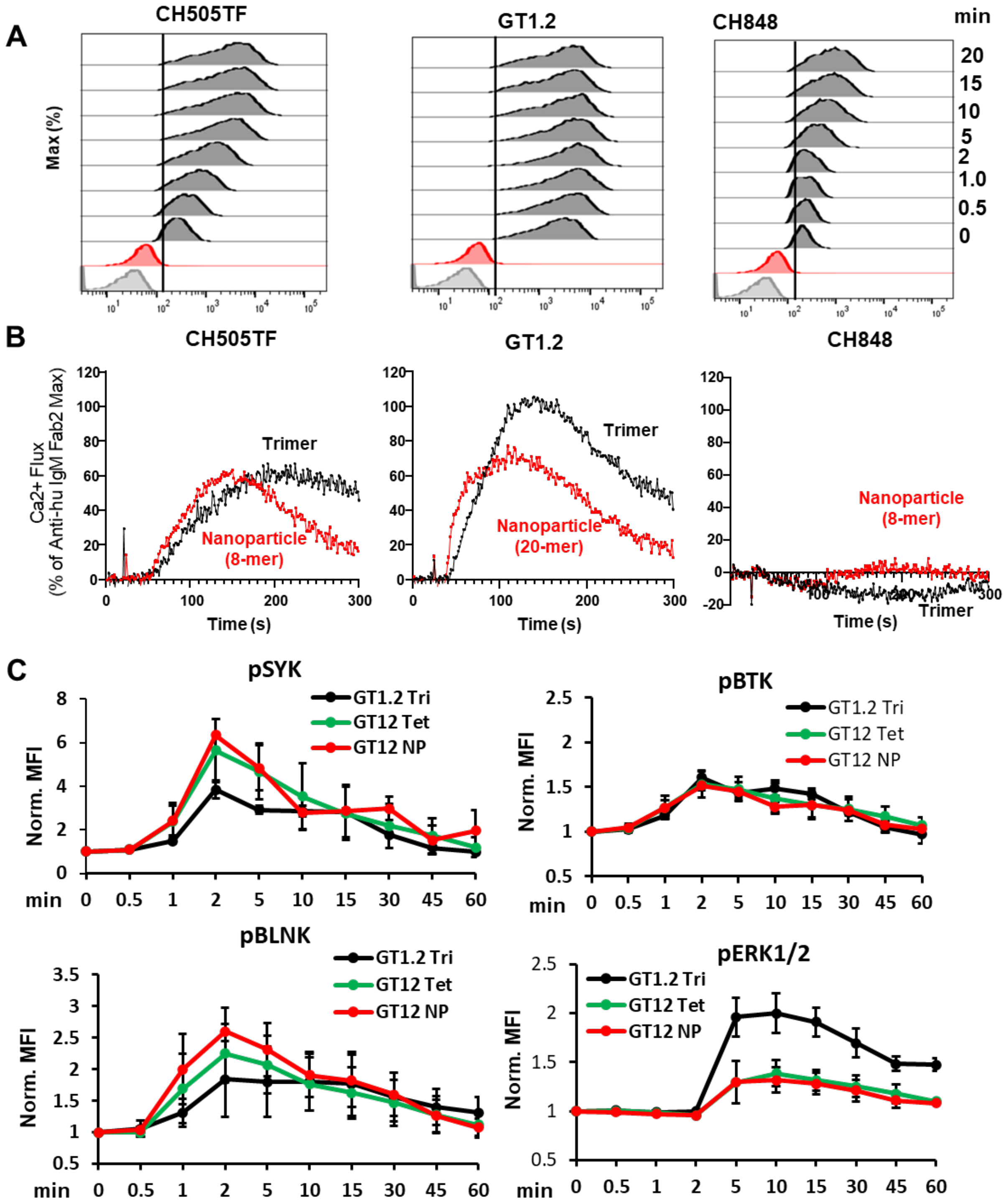


**Figure S6: Cell surface binding and signaling induced by SOSIP trimers and multimers of trimers.** (**A**) Representative flow plots showing binding of CH505TF, GT1.2 and CH848 trimers to CH31 IgM BCRs over a period of 20 mins. Cells were stimulated with the biotinylated trimeric antigens at 30 nM concentration at 37$^{o}$C and aliquots of cells were collected at the indicated timepoints which further fixed and stained by anti-biotin antibody to analyzed in flow cytometer. (**B**) Comparison of the CH31 IgM

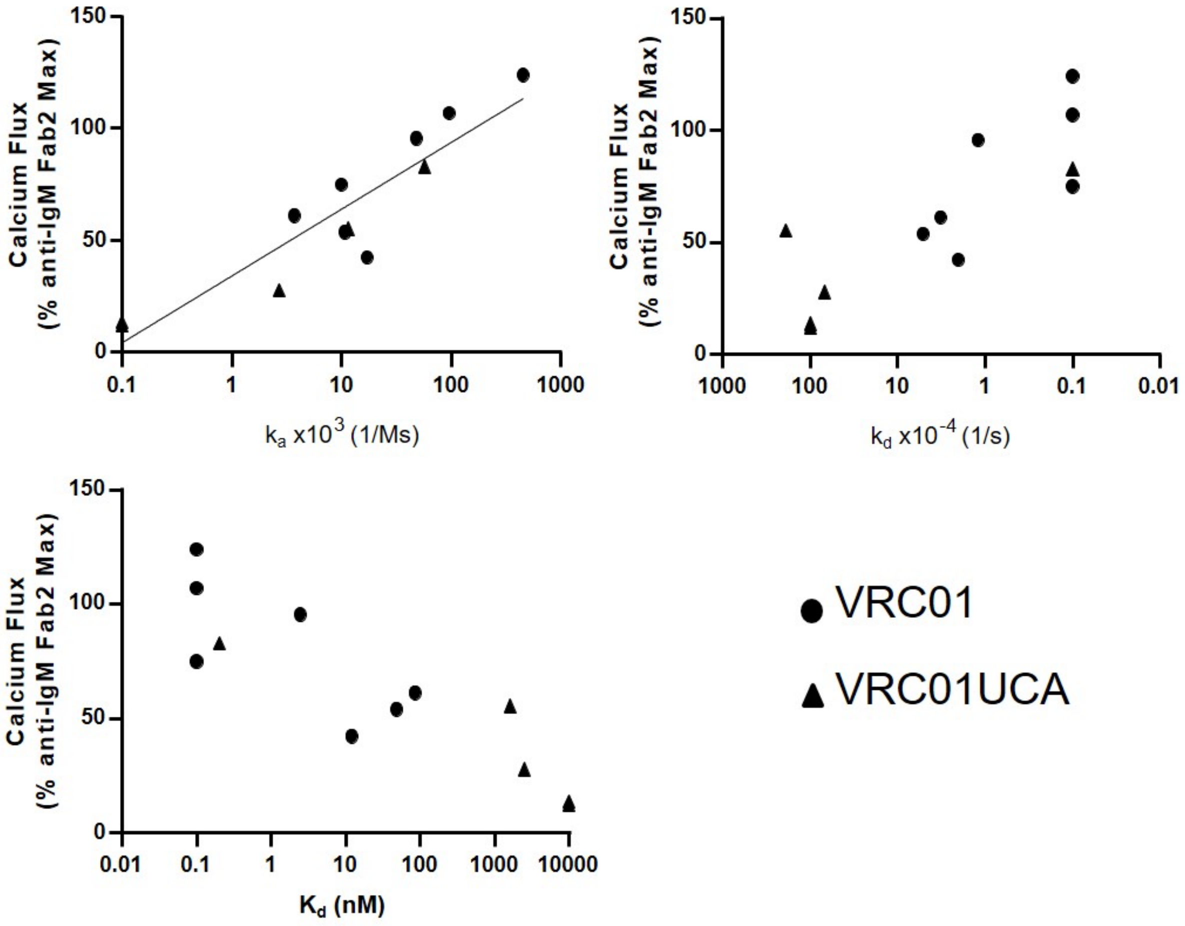


**Figure S7. Relationship between calcium flux response and antigen binding kinetic rates for VRC01 and VRC01UCA Ramos cell lines**. VRC01 and VRC01UCA IgM Ramos cell calcium flux mediated by multimerized antigens (y-axis) plotted versus SPR measured monomeric (or trimeric) antigen on-rate ($k_a$, 1/Ms) (top left), off-rate ($k_d$, 1/s) (top right) or affinity ($K_d$, nM) (bottom left) against VRC01 or VRC01UCA bnAb (x-axis). Calcium flux responses are presented as a percentage of the maximum Anti-human IgM F(ab')2 mediated responses and both calcium flux and kinetic rate parameters are representative one measurement for VRC01/VRC01UCA results. Correlation evaluation was assessed by Kendall's Tau analysis ($k_a$ and VRC01 UCA: Kendall's Tau 0.9487; p-value = 0.0230; $K_d$ and VRC01 UCA: Kendall's Tau -0.9487; p-value = 0.0230). The association between $K_d$ and VRCO1UCA IgM Ca-flux did not reach significance ($K_d$ and VRC01: Kendall's Tau -0.6172; p-value = 0.0599).

**Methods:** Calcium flux experiments were performed as described in the methods section "Calcium Flux Measurements with Ramos cells". SPR affinities and kinetic rates were measured as previously described (Bonsignori, et al., 2018). To assess potential correlations, Kendall's Tau was used due to the small sample size. The alpha level was set at 0.05 and there were no adjustments made for multiple correlation evaluations. The analysis was conducted using SAS 9.4 software.

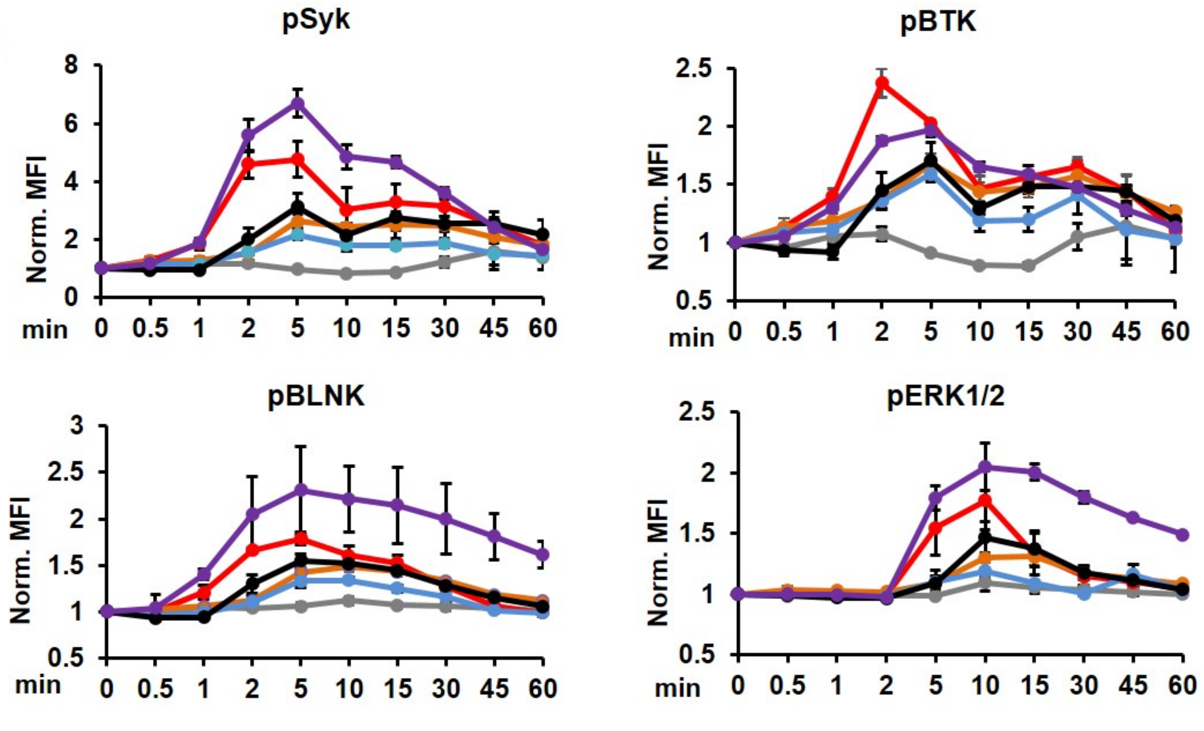


**Figure S8. CH31 IgM B phospho signaling by tetrameric antigens.** B cell signaling kinetics by tetramers of gp120 based antigen stimulation. The kinetics of proximal Syk, BLNK, Btk and ERK1/2 phosphorylation in CH31 IgM Ramos cells were measured over an hour of stimulation by GT6 (grey), GT8 (red), A244 (orange), CH505TF (blue) and 426c (black) at 250 nM concentration along with anti-human IgM (purple) at 30 nM concentration. Around 2.5x10$^5$ cells were collected at each indicated period of times, fixed (30 mins), permeabilized and then stained for phosphorylated Syk, BLNK, Btk and ERK1/2. Plots shown here is normalized MFI data from three independent experiments. At each time point, the phosphorylation level was normalized to the unstimulated cells MFI value (normalized value =1 for the unstimulated, unphosphorylated subset). Error bars indicate standard deviation of means.

**Methods:** Experimental conditions are as described in the methods section under “Antigen binding analysis in CH31 IgM Ramos cells” and “Phospho-flow kinase analysis in CH31 IgM Ramos”.

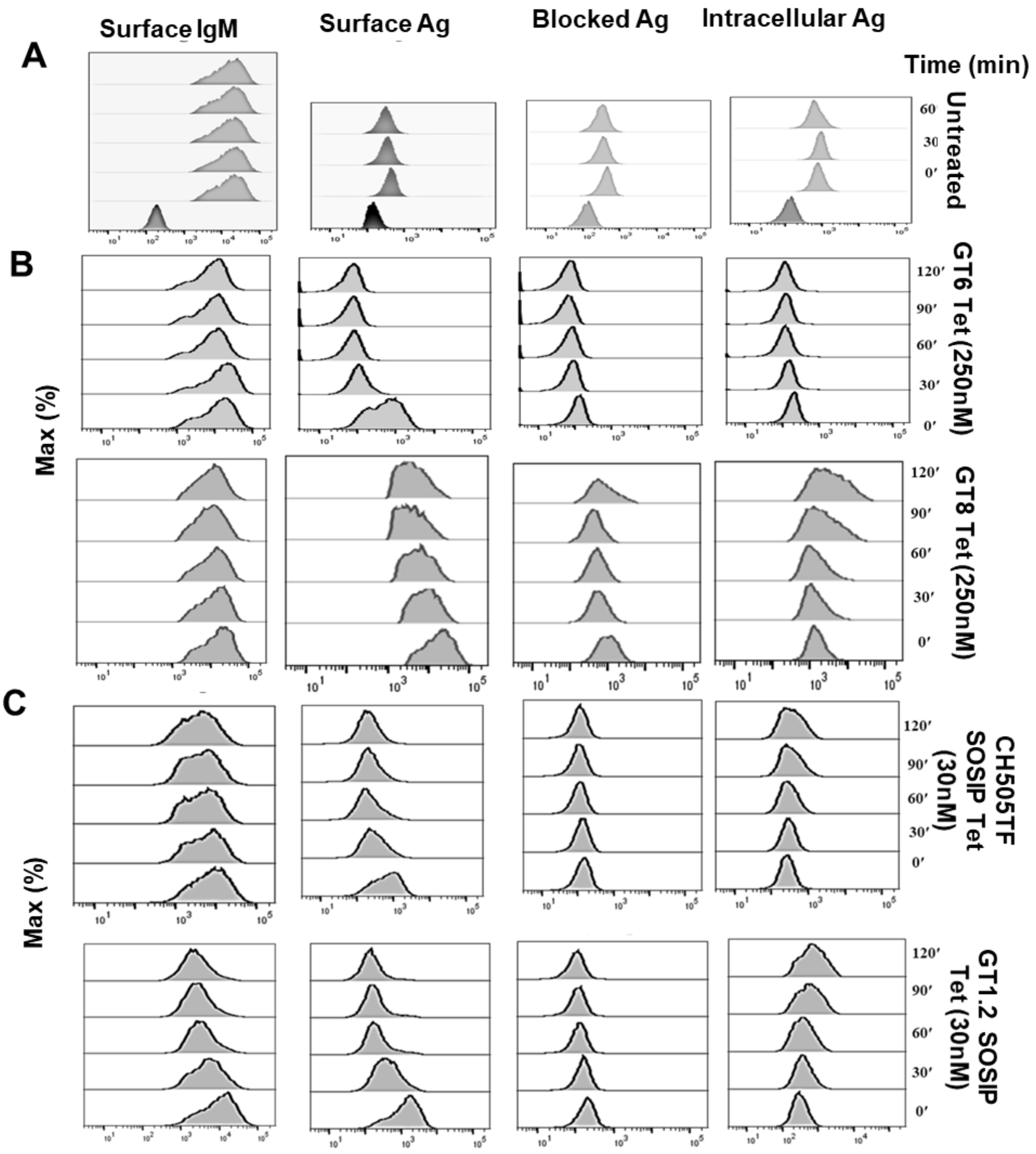


**Figure S9. Flow cytometric detection of intracellular antigens in CH31 IgM Ramos cells**. Representative flow plots showing level of surface IgM, surface antigens, blocked antigens and intracellular antigens detected over time in ramos cells incubated with SA-conjugated tetramers of gp120 based antigens **(B)** and tetramer of trimeric antigens **(C)** or left untreated **(A)**. Level of surface IgM and bound antigens were detected at each time points using flourophore conjugated anti-IgM and anti-SA antibody respectively. To block the SA-binding sites on surface antigens, cells were incubated with non-fluorophore anti-SA antibody and blocked level of SA binding sites were confirmed. For intracellular antigen detection, cells were fixed and permeabilized after surface blockage of SA binding sites and further incubated with fluorophore conjugated anti-SA antibody and analyzed by flow cytometry.

**Methods:** Experiments were performed as described in the methods section under "Antigen internalization assay in CH31 IgM Ramos".

| CH31 IgG mAb | | | | |
|---|---|---|---|---|
| Ag | Assay Temp (°C) | Monomer/Trimer (RU) | Tetramer (RU) | Response (RU) Fold Change |
| 426c degly3 | 25 | 7.8 | 105.9 | 13.6 |
| GT6 | 25 | 24.2 | 95.6 | 4.0 |
| GT8 | 25 | 69.5 | 137.8 | 2.0 |
| CH505TF | 25 | 65.2 | 98.6 | 1.5 |
| 426c | 25 | 62.5 | 278.7 | 4.5 |
| A244 | 25 | 57.7 | 51.0 | 0.9 |
| CH505TF Trimer | 37 | 243.6 | 276.0 | 1.1 |
| GT1.2 Trimer | 37 | 570.4 | 593.5 | 1.0 |

**Figure S10. Response fold change of antigen monomers/trimers versus multimers.** Summary table comparing the SPR binding responses (RU) to CH31 IgG bnAb of monomeric antigens (or trimeric) with the binding responses of their multimeric (4-mer) forms. The responses were measured at the end of the association phase of the sample injection and the fold change between the antigen types was calculated by dividing the response of the tetramer by the response of the monomer (or trimer). The 426c degly3 and 426c core proteins exhibited the largest increase in binding response against CH31 IgG after tetramerization. GT6, GT8, and CH505TF gp120 all showed some enhancement in binding (1.5x or greater) after tetramerization while A244, CH505TF trimer, and GT1.2 trimer showed little to no increase in binding.

**Methods:** SPR experiments with monomeric (or trimeric) antigens versus low-order multimers of the same antigens were performed as described in the methods section under "Surface Plasmon Resonance of Tetrameric Antigens".
